# Stimulation of the Medial SNr Promotes Sustained Motor Recovery and Counteracts Parkinsonian Pathophysiology in Dopamine Depleted Mice

**DOI:** 10.1101/2024.12.09.627637

**Authors:** Asier Aristieta, John E. Parker, Mary D. Cundiff, Thomas Fuchs, Byung Kook Lim, Jonathan E. Rubin, Aryn H. Gittis

## Abstract

Dopamine loss alters the activity of neural circuits in the basal ganglia, contributing to motor symptoms of Parkinson’s disease and catalepsy. Treatments that reduce basal ganglia pathophysiology alleviate motor symptoms but require maintenance. Cell-type specific interventions can reduce pathophysiology and provide sustained therapeutic benefits, but a lack of understanding of pathways involved limits translation. Here, we establish patterns of neuromodulation and electrophysiological biomarkers at the level of basal ganglia output that predict the duration of therapeutic effects. Focal activation of neurons in the ventromedial substantia nigra reticulata (SNr) engaged a gradual recovery of movement that persisted for hours after treatment, accompanied by a persistent reduction in parkinsonian pathophysiology. Global SNr inhibition, as prescribed by the classic rate model, provided only transient effects on movement and did not reverse network pathophysiology. These findings represent important steps towards developing therapeutic strategies that aim to repair, rather than simply mask, circuit dysfunction in disease.

## Introduction

Impaired dopamine signaling results in decreased motor output associated with conditions such as Parkinson’s disease and catalepsy. The basal ganglia are the main recipient of dopamine released from the substantia nigra pars compacta (SNc). The loss of SNc dopamine neurons causes reduced motor output that is often accompanied by changes in neural activity throughout the basal ganglia, culminating in disruption of both firing rates and firing patterns of basal ganglia output neurons^1–6^. As a major basal ganglia output nucleus, the substantia nigra pars reticulata (SNr) is poised to propagate pathological activity to dozens of target structures across the brain, disrupting brain-wide motor pathways required for movement^7,8^.

Treatments such as deep brain stimulation (DBS) and pallidotomy have established that mitigating basal ganglia output pathophysiology can promote movement under conditions of low dopamine^9–15^. Although preferable to irreversible lesions of basal ganglia output, DBS requires constant stimulation to maintain its effects. When stimulation is turned off, motor symptoms return within minutes^16^. This transient efficacy of DBS suggests that it works by masking basal ganglia output pathophysiology, rather than by restoring underlying circuit function^9,17,18^.

The goal of next generation therapies for movement disorders will be to transition from palliative care to treatments that can restore circuit function and provide sustained therapeutic benefits. Towards this goal, previous work from our lab found that long-lasting locomotor recovery can be induced by targeting neuronal subpopulations in the external segment of the globus pallidus (GPe)^19,20^. Interventions that transiently increased the activity of parvalbumin-expressing neurons (PV-GPe) or decreased the activity of lim-homeobox 6-expressing neurons (Lhx6-GPe) induced sustained recovery of movement that persisted for hours beyond the period of active stimulation. But a lack of understanding of the neural circuits mediating these long-lasting therapeutic effects remains a hurdle for translation.

The SNr is a major target of both PV-GPe and Lhx6-GPe subpopulations and interventions that induce persistent motor rescue share a common mechanism of reducing burst firing in the SNr, a feature of parkinsonian pathophysiology^19,21^. This result would seem to implicate the SNr as a key site for the induction of persistent rescue, but stimulation of direct pathway neurons in the striatum, whose main target is also the SNr, induces only acute locomotor effects that are not persistent^19^. Additionally, if both PV-GPe and Lhx6-GPe subpopulations inhibit the SNr, why must they be oppositely modulated in order to induce persistent rescue? These questions prompted an investigation into the patterns of activity locally within the SNr that are required to convert short-term to long-term motor recovery.

We find that persistent therapeutic effects are induced by activation of a subset of ventromedial SNr neurons located in the region receiving input from Lhx6-GPe neurons. Optogenetic stimulation of these ventromedial SNr neurons was sufficient to drive a gradual recovery of movement that accumulated over the course of the ∼30 minute treatment period and persisted for several hours after stimulation. Conversely, global inhibition of the SNr, as prescribed by the classic rate mode^22,23^, drove acute motor responses that did not persist outside of the stimulation period. Intriguingly, stimulation of the medial SNr attenuated some features of parkinsonian pathophysiology and thereby transitioned the nucleus into a healthier physiological state that persisted for several hours after stimulation. These results provide a physiological template for the induction of sustained motor recovery under conditions of low dopamine.

## Results

### SNr Lesions Are Protective Against Parkinsonian Motor Deficits

Abnormal activity of the basal ganglia is thought to be a primary contributor to motor dysfunction following dopamine loss^1–6^. Consistent with this idea, surgical ablation of the internal globus pallidus (GPi) has been used to treat motor symptoms of Parkinson’s disease in patients and in non-human primate models of the disease^24^. In non-primate mammals, the SNr, not the GPi, is the main basal ganglia output nucleus^25^. Aberrant neural activity in the SNr has been shown to correlate with the onset and progression of motor symptoms in rodent models of dopamine depletion^5,26,27^, but whether SNr dysfunction is causal to motor symptoms has not been explicitly tested.

To determine the contributions of the SNr to the parkinsonian phenotype observed in the bilateral 6-hydroxydopamine (6-OHDA) dopamine depletion model^28^, we assessed the effects of dopamine depletion on motor function in mice with or without an intact SNr. Cellular ablation of the SNr was achieved using a Cre-targeted diphtheriatoxin strategy^29^: AAV carrying a Cre-inducible diphtheriatoxin receptor (AAV2-DIO-DTR-GFP) was stereotaxically injected into the bilateral SNr of PV-Cre mice, followed 3 weeks later by an IP injection of 1 µg diphtheriatoxin (DTX) (Fig. 1a). Approximately 90% of SNr neurons have been shown to exhibit some degree of PV expression^7^, which we confirmed in our Pvalb-2A-Cre line^30^ with injections of the color changing reporter virus Nuc-flox(mCherry)-EGFP. Cre-mediated color conversion was observed in 86% of SNr neurons (Fig. 1b). Successful ablation of the SNr was confirmed postmortem by a reduction in DTR-GFP fluorescence signal in the SNr of mice treated with DTX compared to those who had received saline (Fig. 1c,g). Two weeks after administering DTX to ablate the SNr (or saline injections for SNr-intact controls), mice were dopamine depleted with bilateral 6-OHDA injections into the medial forebrain bundle. Dopamine depletions were confirmed postmortem with TH immunoreactivity in the striatum compared to littermate controls (Fig. 1f).

**Figure 1:**
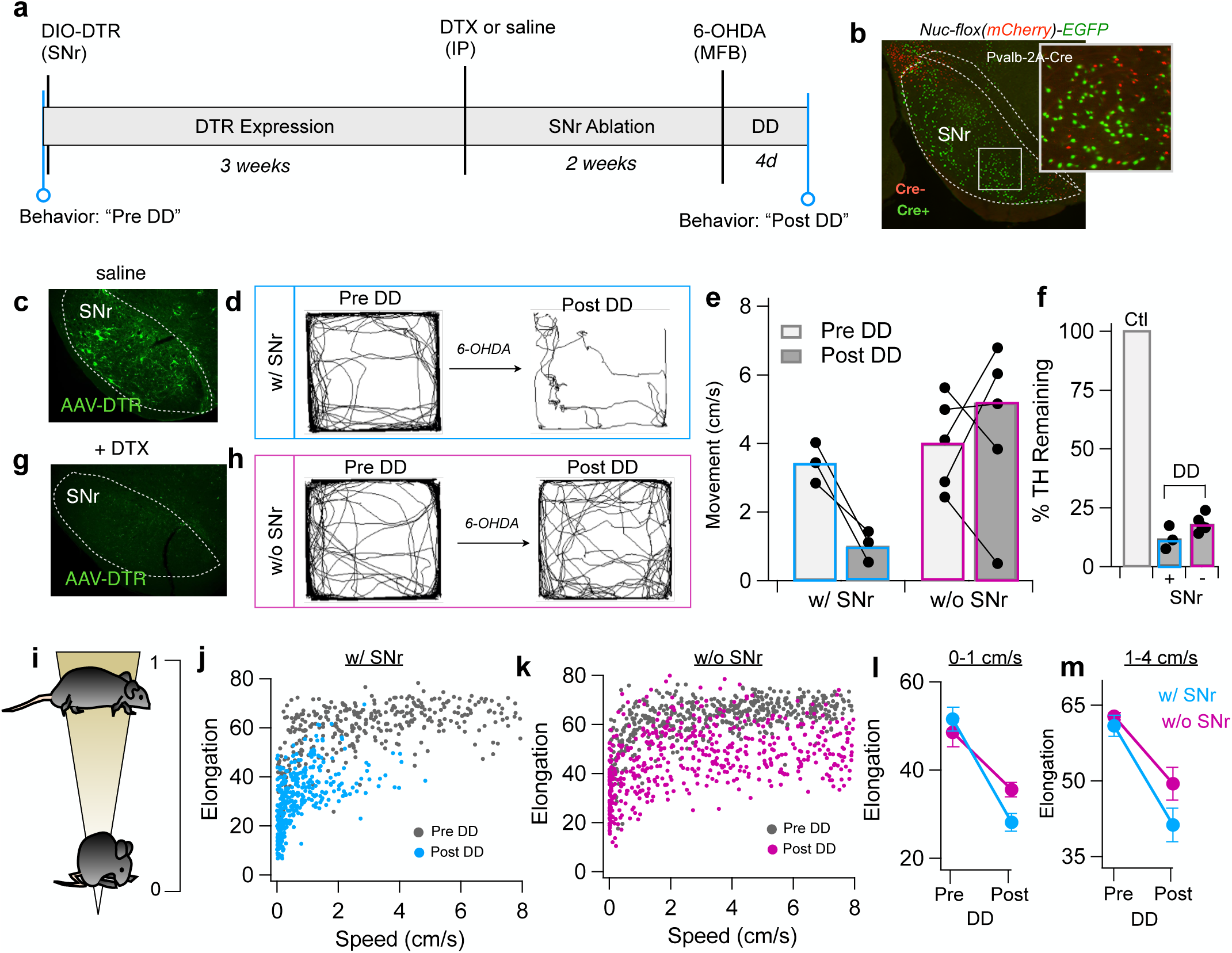
Ablation of SNr neurons is protective against locomotor impairments in dopamine depleted mice. **a,** Experimental timeline. **b,** Fluorescent image of the SNr from a PV-Cre mouse injected with AAV-Nuc-flox(mCherry)-EGFP showing the distribution of Cre^+^ cells (green) throughout the nucleus. Cre^-^ cells are red. **c,g,** Fluorescent signal in the SNr showing the distribution of neurons expressing DTR-GFP from an SNr-intact control subject (IP saline, *c*) or an SNr-ablated subject (IP DTX, *g*). **d,h,** Movement paths over a 20 min period recorded in representative mice that were SNr-intact (*d*) or SNr-ablated (*h*) before and after bilateral dopamine depletion. **e,** Group average showing movement speed (cm/s) recorded in the open field before vs. after bilateral dopamine depletions in SNr-intact mice or SNr-ablated mice. Paired pre/post data points show data from individual animals. Paired t-test: SNr intact (N = 3), p = 0.04; SNr-ablated (N = 5), p = 0.696. **f,** TH immunoreactivity measured in the striatum of dopamine depleted mice, normalized to littermate controls. Paired t-test: SNr-intact (N = 3) vs SNr-ablated (N = 5), p = 0.09. **i,** Schematic of postural quantification. **j-k,** Body elongation (5 s bins) plotted as a function of movement speed for dopamine depleted mice in the SNr-intact (blue, *j*) or SNr-ablated (magenta, *k*) groups. Grey dots are data from dopamine-intact littermates (N = 2, same data plotted in both *j* and *k*). **l-m,** Change in body elongation score pre vs. post dopamine depletion for mice in the SNr-intact (blue, -N = 3) and SNr-ablated (magenta, N = 5) groups. At low movement speeds: (*l*, 0-1 cm/s): 2-way ANOVA showed a significant main effect of dopamine depletion (F(1,12) = 39.7, p < 0.0001), and no significant interaction between SNr groups (F(1,12) = 3.67, p = 0.079). At high movement speeds: (*m*, 1-4 cm/s): 2-way ANOVA showed a significant main effect of dopamine depletion (F(1,12) = 33.4, p < 0.0001), and no significant interaction between SNr groups (F(1,12) = 1.85, p = 0.199).

In SNr intact mice, bilateral dopamine depletions strongly suppressed movement, typical of this depletion model. Movement in an open field arena was reduced from an average of 3.4 ± 0.6 cm/s before depletion to 1.0 ± 0.5 cm/s after depletion (p = 0.04, N = 3) (Fig. 1d,e). In contrast, mice whose SNr had been previously ablated using DTX were relatively spared from the motor impairing effects of dopamine depletion. Mice freely explored the open field arena and moved to similar degrees after depletion (4.5 ± 1.6 cm/s) as before (4.0 ± 0.6 cm/s, p = 0.70, N = 5) (Fig. 1e,h).

Given the protective effects of SNr ablation on movement, we sought additional confirmation, beyond histology, that dopamine depletions had been successful. In addition to decreasing movement, bilateral dopamine depletions cause postural changes characterized by a hunched or stooped posture that persists even during walking (Fig. 1i). The neural circuits underlying postural changes in Parkinson’s disease remain obscure, but because postural symptoms are rarely responsive to dopaminergic treatment (e.g. L-DOPA)^31^, they have been hypothesized to involve circuits outside of the basal ganglia^32,33^.

To quantify postural changes induced by dopamine depletion in our mice, we used an elongation metric (Noldus) that reports posture as a percentage of a mouse’s fully extended body length: low values indicate contracted postures and high values indicate stretched postures (Fig. 1i). As shown in Fig. 1j-k, mice are more contracted at rest and elongate during locomotion. In both groups of mice (SNr intact and SNr lesioned), dopamine depletion caused contraction of body posture, resulting in a downward shift in the body elongation metric across all speeds measured (Fig. 1j-m). At low speeds/rest (0-1 cm/s), dopamine depletion drove a significant contraction of body posture in both SNr-intact and SNr-lesioned mice (2-way ANOVA, F(1,12) = 39.7, p < 0.0001); the effect size was not significantly different between SNr lesioned vs. intact groups (p = 0.43) (Fig. 1l). Postural deficits persisted in both groups during locomotion (Fig. 1m). There was a significant main effect of dopamine depletion on body posture (2-way ANOVA, F(1,12) = 33.4, p < 0.0001) but no main effect of SNr group (intact vs. lesioned, p = 0.06) (Fig. 1m).

Taken together, these results suggest that aberrant output from the SNr contributes to decreases in movement following bilateral dopamine depletion, but not to changes in posture.

### Identifying Predictive Biomarkers of SNr Pathophysiology Using a Neural Network Classifier

Under dopamine depleted conditions, changes in firing rates and patterns of SNr neurons have been reported, but there is no clear consensus on which features of electrophysiological activity are most indicative of a physiologically “normal” or “pathological” state. SNr neurons may exhibit increases or decreases in firing rates after dopamine depletion and aberrant firing patterns have been described in terms of bursts, synchrony, oscillations, and irregularity^4,5,26,34–38^. Additionally, because SNr pathophysiology is typically described in terms of population metrics (group averages before vs. after depletion), specific biomarkers for tracking pathophysiology at the level of individual neurons across treatment periods are lacking.

To develop a rigorous quantitative method to assess SNr pathophysiology at the level of individual neurons, we conducted a meta-analysis of a large physiological data set that included n = 512 SNr neurons recorded across N = 14 dopamine intact mice and n = 623 SNr neurons recorded across N = 26 dopamine depleted mice (Fig. 2a).

**Figure 2:**
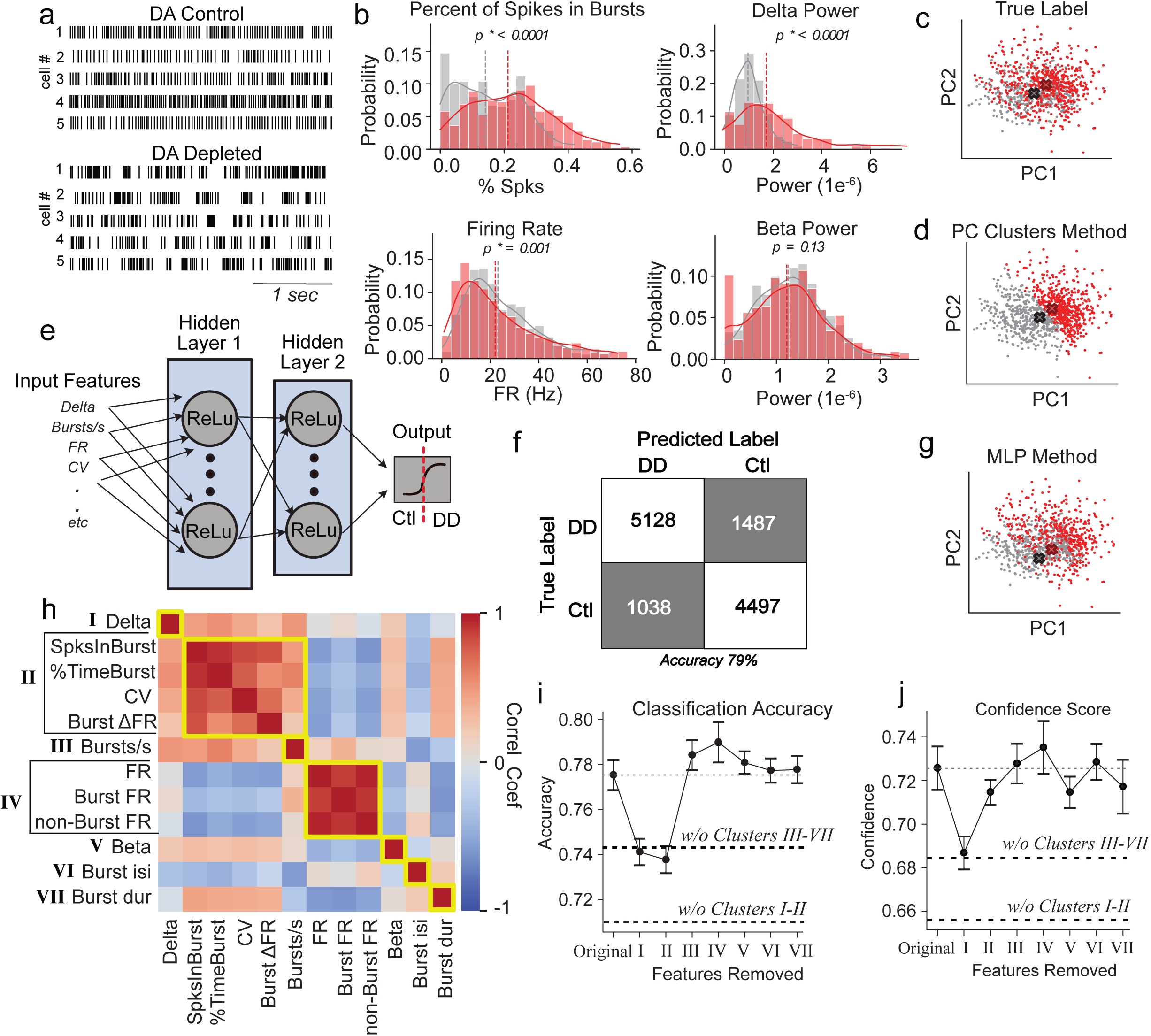
A machine learning approach for quantifying pathophysiological severity at the level of individual neurons. **a,** Rasters showing spontaneous activity of representative SNr neurons under dopamine-intact control (*top*) or dopamine depleted (*bottom*) conditions. **b,** Histograms of the values observed for individual electrophysiological features recorded from dopamine-intact control mice (grey, n = 512 neurons, N = 14 mice) and dopamine depleted mice (red, n = 623 neurons, N = 26 mice). The Kolmogorov-Smirnov test revealed a significant difference between control vs. depleted groups in percentage of spikes in bursts (p < 0.0001), delta power (p < 0.0001), firing rate (p < 0.001), but not beta power (p = 0.13). **c,** PCA feature space (PC1 vs PC2) projections of electrophysiological features measured from recordings in control (grey) and dopamine depleted (red) animals. **d,** Same data as *c*, but with colors assigned based on the nearest centroid computed using the L^2^ norm. Centroids were determined using the k-means clustering algorithm and projected into PC1 vs PC2 space. **e,** Schematic of multi-layer perceptron (MLP) artificial neural network: an input layer, two hidden layers with 200 and 100 units respectively, and an output layer. All layers used the ReLU activation function and were connected in a feed-forward manner. **f,** Cumulative confusion matrix showing classification results on training data for 15 seeds of the MLP. Reported accuracy 79.2 ± 0.83% is averaged over all 15 models. **g,** Same data as *c*, but with colors assigned based on classification by averages of probability scores from all 15 MLP models. **h,** Correlation matrix showing relationships between the 12 electrophysiological features used for classification. Scale bar is correlation coefficient. Clusters II and IV are determined by parameters correlated (C>0.8) with, respectively, %SpksBurst (Cluster II) and firing rate (Cluster IV). **i,** Average accuracy of trained MLP models applied to all data was 77.5 ± 1.3%. When features in Cluster I and II were withheld, model accuracy dropped to 71%. When features in Clusters III-VII were withheld, model accuracy was 74%. **j,** MLP classification DD confidence score assigned when various features were withheld from the model. When features in Cluster I and II were withheld, model confidence in DD classification dropped to 66%. When features in Clusters III-VII were withheld, model DD confidence was 68%.

A collection of 12 spike train features, including firing rate, CV, power in delta and beta frequency bands, and various bursting indicators (see Methods for detailed definitions), were analyzed to determine which neural firing properties best differentiated between dopamine intact and dopamine depleted conditions. Numerous features were significantly different at the population level between control vs. dopamine depleted data sets, a subset of which are shown in Fig. 2b. On average, spike trains recorded from depleted mice showed more bursting activity than those recorded from controls, increased power in the delta frequency band and a small but significant decrease in firing rates (Fig. 2b). Inspection of beta power in the spike trains revealed no significant decrease between control and depleted mice (Fig. 2b), as previously reported in mice^26,39^. As evident from the highly overlapping distributions seen in all these histograms, it is not possible to use any single parameter to identify a neuron’s level of pathophysiology.

To establish a multi-parametric classification strategy, we first attempted to use PCA on the feature space (Fig. 2c-d). Although clustering of neurons in PC space was able to distinguish depleted from control populations with greater accuracy than single parameter thresholds, separation was still poor (Accuracy = 64%). Consequently, we employed a multi-layer perceptron (MLP) artificial neural network, which is a supervised machine learning algorithm that is well-suited for classifying and modeling complex data. The MLP model consisted of an input layer for the 12 features of the spike train recordings, followed by two hidden layers that applied nonlinear transformations of the data, culminating in an output layer that classified an input spike train as either recorded from a control or dopamine depleted mouse (Fig. 2e). We used a randomly selected 80% of our baseline healthy data and 80% of our baseline depleted data to train the MLP and then tested its performance on the remaining 20% of the data. We repeated this process de novo fifteen times with independent selections of training data. Across these simulations, the MLP achieved an average accuracy of 79.2 ± 0.83% on the training data (Fig. 2f) and 70.8 ± 0.8% on the test data. When projected to PC space, the clusters identified by the MLP showed notable mixing, as seen with the correctly labeled points in PC space determined by the average of the trained MLP models’ confidence scores (Fig. 2c,g).

The MLP relied on a combination of all features to distinguish between the dopamine intact and dopamine depleted states (Fig. 2h). Among these, power in the delta frequency band (0.5-4 Hz) emerged as a prominent feature used by the model to distinguish between control and depleted states. Removal of this feature decreased the model’s classification accuracy and confidence scores for identifying neurons recorded from depleted animals (Fig. 2i-j). Several features related to firing rate irregularity were highly correlated with each other (“Cluster II”): elevations in the percentages of spikes occurring within bursts (SpksInBurst), the percentage of time spent bursting (%TimeBurst), the increase in firing rate within bursts (BurstΔFR), and firing CV (Fig. 2h). Removing these features from the model decreased accuracy to a similar degree as removing delta power (Fig. 2i), but confidence was not as strongly affected (Fig. 2j). Notably, firing rate itself was not a crucial parameter for differentiating cells recorded from dopamine depleted mice, nor was the number of bursts (Bursts/s) (Fig 2i-j); however, they did provide some information, as evidenced by a decrease in classification accuracy and confidence when these and other parameters were all omitted (Fig. 2i-j, without Clusters III-VII).

### Global SNr Inhibition Promotes Acute Behavioral Rescue

Conventional strategies to treat parkinsonian motor symptoms attempt to reduce inhibitory output of the basal ganglia^1,22,23^. Typically, this is achieved by increasing the activity of D1-expressing spiny neurons in the striatum (D1-SPNs)^22,40,41^, or by direct suppression of basal ganglia output through lesions or pharmacological silencing^4,42–44^. Previously, we have shown that D1-SPN stimulation is sufficient to overcome even extreme parkinsonism associated with bilateral dopamine depletions in the bilateral 6-OHDA model^19^.

Axons of D1-SPNs broadly innervate the SNr (Fig. 3a-b) and drive broad suppression of SNr activity^45^, so it is reasonable to assume that the therapeutic effects of D1-SPN stimulation are mediated by inhibition of the SNr. To test this hypothesis, we measured the motor effects of directly inhibiting SNr activity in bilaterally depleted mice (N = 6). To globally inhibit neural activity in the SNr, we injected a pan-neuronal inhibitory opsin (hSyn-Jaws) into the bilateral SNr (Fig. 3a,c). After allowing 3 weeks for expression, mice were bilaterally dopamine depleted and behavioral experiments were performed 4-5 days later.

**Figure 3:**
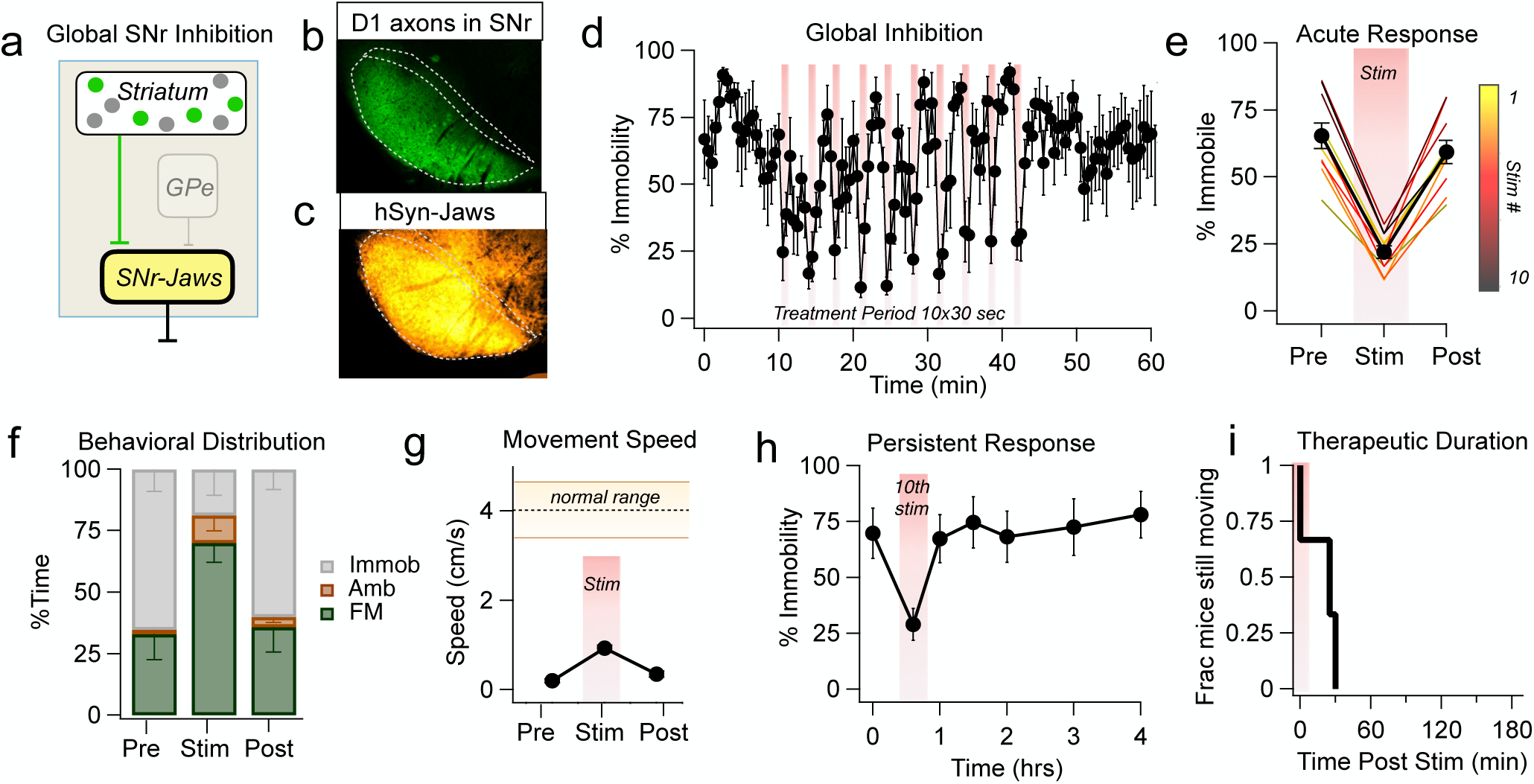
Global inhibition of the SNr promotes movement acutely. **a,** Experimental schematic. **b,** Fluorescent image of D1-SPN axons in the SNr. **c,** Fluorescent signal (pseudo-color heat map) in the SNr, averaged across N = 6 mice expressing hSyn-Jaws-mCherry. Section alignment and digital averaging were performed with ImageJ. **d,** Immobility (percentage of time mice exhibited >0.01% pixel change in 30 s bins) as a function of time during treatment with global inhibition, N = 6 mice, error bars, sem. Red lines denote periods of optical stimulation of Jaws: 30 s red light (1 mW, continuous), spaced 3 min apart. **e,** Acute response to Jaws activation, averaged across all mice. Averages shown for each trial (1-10, thin lines) and across all trials (black circles, error bars, sem). Data are from 30 s immediately preceding each stimulation (*pre*), 30 s during stimulation (*stim*), and from 30 s – 60 s after each stimulation (*post*). Immobility was significantly decreased during the lightON period. Paired t-test: Pre vs. Stim (N = 6 mice) p = 0.0009. **f,** Fraction of time mice spent immobile (*grey*) or engaged in fine movements (*green*) or ambulation (*brown*). **g,** Average movement speed calculated pre, during, and post each light stimulation. Average movement speed during lightON periods was 0.92 ± 0.23 cm/s, N = 6 mice. Error bars, sem. The average movement speed of dopamine intact controls was 0 ± 1.4 cm/s (shaded region, ± sem, N = 5). **i,** Therapeutic duration, calculated from the time of last stimulation until mice returned to within 80% of their initial immobility.

In an open field arena (50 x 50 cm^2^), dopamine depleted mice spent the majority of their time in the immobile state (Fig. 3d). Optogenetic inhibition of the SNr (1 mW continuous stim for 30 s, repeated every 3 min) acutely reduced immobility during the lightON periods, with no carry-over effects observed in between or after the conclusion of light pulses (Fig. 3d-e).

To examine the nature of movements that occurred during SNr inhibition, behaviors were classified as “fine movement” (stationary movements such as grooming, sniffing, scratching), “ambulation” (locomotor behaviors such as walking, running), or “immobile” (no movement). Using this analysis, we found that the decrease in immobility driven by SNr inhibition was largely due to an increase in fine movements (Fig. 3f). During the lightON periods, movement increased from 0.20 ± 0.15 cm/s to 0.92 ± 0.23 cm/s, but this was still far below the amount of movement seen in dopamine intact mice (3.8 ± 1.1 cm/s) (Fig. 3g).

After the treatment period, mice were kept in the open field for an additional 3 hrs to test the persistence of movement effects. As shown in Fig. 3h-i, therapeutic effects were not persistent, and mice returned to pre-treatment levels of immobility within minutes (average = 12.5 min; range = 0-30 min).

### Global SNr Inhibition Does Not Reduce Parkinsonian Pathophysiology

The pro-kinetic effects of global SNr inhibition are consistent with the concept that reducing basal ganglia output is therapeutic under conditions of low dopamine, but the effects are transient. To assess the impact of global inhibition on SNr pathophysiology, we performed recordings from awake head fixed mice. An optrode consisting of a 16-site linear array and integrated optical fiber was lowered into the SNr to record neural activity during optogenetic treatment (1 mW continuous stim for 30 s, repeated every 3 min). In mice expressing AAV-hSyn-Jaws in the SNr, optogenetic stimulation drove robust suppression of firing in 26 out of 27 neurons (Fig. 4a-b).

**Figure 4:**
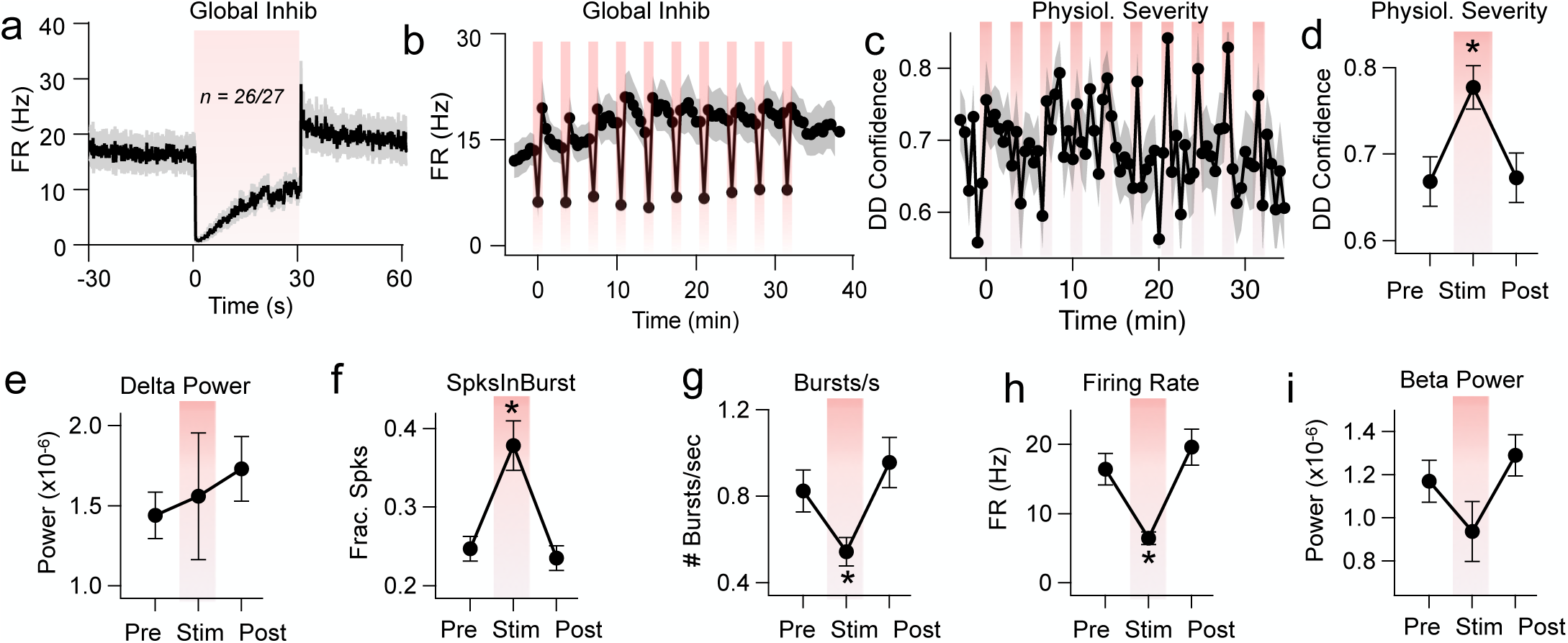
Electrophysiological changes induced by global SNr inhibition are transient. **a,** Responses of SNr neurons during 30 s light activations of Jaws (1 mW, continuous red light, 10 repetitions spaced 3 min apart). Of the 27 neurons where recordings were stable across all 10 stimuli (N = 6 mice), 26 were inhibited (FR_pre_ 16.4 ± 10.7 Hz vs. FR_stim_ 6.5 ± 4.2 Hz; paired t-test, p = 0.0003). The only neuron that was not inhibited showed no significant firing rate modulation during light stimulation. **b,** Firing rates were reliably inhibited across all 10 stimuli. **c,** MLP confidence scores in 30 s bins (average of n = 27 neurons, error bars sem) indicating the model’s confidence that a given neuron was from a dopamine depleted animal (chance = 0.5). **d,** MLP confidence that neurons exhibited parkinsonian pathophysiology increased significantly during lightON periods: Pre: 0.67 ± 0.13 vs. Stim: 0.78 ± 0.12; paired t-test: p = 0.01). **e-i,** Individual electrophysiological features recorded 30 s before, during, and after optical stimulation. Error bars, sem. There was a significant increase in the fraction of spikes occurring within bursts (SpksInBurst) (Pre: 0.25 ± 0.07 vs. Stim: 0.38 ± 0.03; paired t-test, p = 0.001); significant decreases in the burst frequency (Bursts/s) (Pre: 0.83 ± 0.46 burst/s vs. Stim: 0.54 ± 0.31 bursts/s; paired t-test, p = 0.004) and firing rate (Pre: 16.4 ± 10.7 Hz vs. Stim: 6.5 ± 4.2 Hz; paired t-test, p = 0.0003), and no significant changes in delta power (Pre: 1.4e^-06^ ± 0.68e^-06^ vs. Stim: 1.6e^-06^ ± 0.19e^-06^; paired t-test, p = 0.76) or beta power (Pre: 1.2e^-06^ ± 0.4e^-06^ vs. Stim: 0.94e^-06^ ± 0.64e^-06^; paired t-test, p = 0.16).

To determine whether the activity of individual SNr neurons looked less pathological during stimulation, we used our MLP model (see Fig. 2) to assign confidence values in 30 s continuous intervals that a given unit was likely recorded from a dopamine depleted animal, where 1 meant a unit was definitely recorded in a depleted animal and 0 means the unit was definitely not recorded from a depleted animal. Optogenetic inhibition surprisingly increased, rather than decreased, the model’s confidence that a unit was recorded from a depleted animal (Fig. 4c-d).

To determine which electrophysiological features might be contributing to this effect, we examined at the impact of optogenetic inhibition on individual electrophysiological features (Fig. 2h-j). From Fig. 2, the two most important features used by the model to differentiate activity from control vs. dopamine depleted mice were (1) delta power and (2) burst features, e.g. the fraction of spikes in a burst. Delta power was not significantly altered during optogenetic inhibition (Fig. 4e) and the fraction of spikes within bursts actually increased, rather than decreased (Fig. 4f). Other features of pathophysiology were decreased during optogenetic inhibition including the number of bursts (Fig. 4g), firing rate (Fig. 4h), and beta power (Fig. 4i), but these parameters were shown to be less important for the discrimination accuracy of the MLP network (Fig. 2i-j). These results indicate that global inhibition reduces some features of SNr pathophysiology but not the features that are the most predictive of the dopamine depleted state.

### Npas1-Cre Line Preferentially Labels Medial SNr Neurons

Previously, we have shown that long-lasting recovery of movement can be induced in dopamine depleted mice by cell-type specific interventions in the GPe^19,20^. The transient inhibition of Lhx6-GPe neurons is crucial to induce a sustained therapeutic response. This requirement is counterintuitive to the classical rate model, which predicts that excitation of the GPe, not inhibition, should be therapeutic. We wondered if the innervation pattern of Lhx6-GPe neurons to the SNr might provide insights into the privileged role of this population for long-lasting motor recovery.

Using traditional AAV-based fluorescent markers, the projections of Lhx6-GPe neurons to the SNr appear similar to those of other GPe neurons (e.g. PV-GPe)^21^. But partial overlap between neurons labeled in the Lhx6-GPe and PV-GPe mouse lines might obscure important anatomical differences^46^. To resolve the projection patterns of Lhx6_only_-GPe neurons that do not express PV, we used an intersectional transgenic approach in which Lhx6-iCre mice were crossed to PV-Flp mice to create Lhx6-iCre;PV-Flp heterozygotes. We then virally transfected the GPe (N = 6 mice) with a fluorescent construct that turns on in the presence of Cre, but turns off in the presence of Flp (rAAV_DJ_-pAAV-Ef1α-FRT-DIO-eGFP) (Fig. 5a). This enabled restricted expression of fluorescent protein in the subset of Lhx6_only_-GPe neurons and visualization of their pattern of innervation to the SNr. As shown in Fig. 5b, axons of Lhx6_only_-GPe neurons were more concentrated in the ventromedial territory of the SNr than other regions. Inhibition of Lhx6-GPe neurons would therefore have a predicted effect of exciting the medial SNr, because GPe neurons are GABAergic.

**Figure 5:**
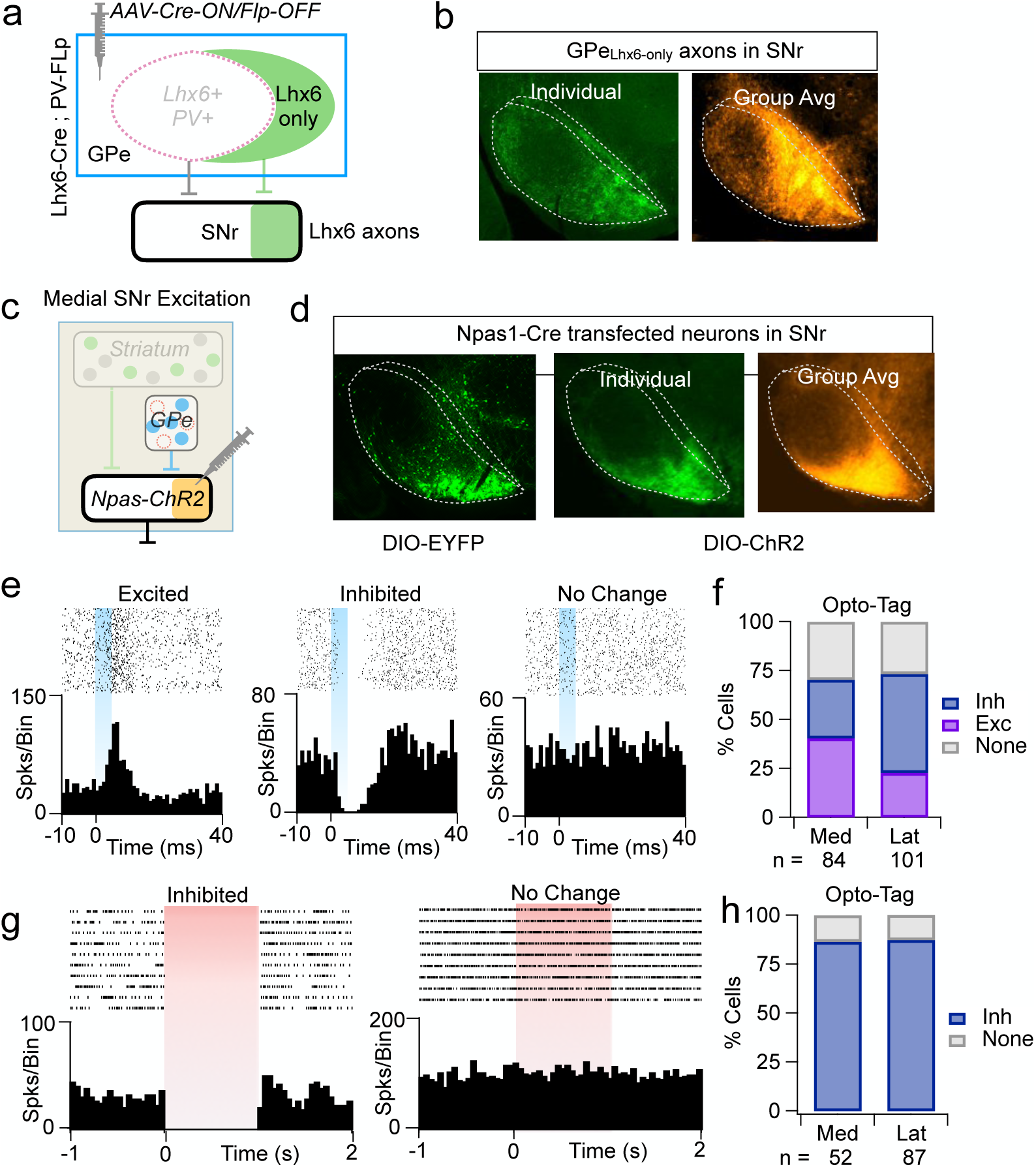
Npas1-Cre line targets neurons in the ventromedial SNr. **a,** Schematic of intersectional strategy to label Lhx6_only_-GPe neurons. **b,** SNr section showing fluorescently labeled axons from Lhx6_only_-GPe neurons (*left*, individual example, green pseudo-color; *right*, average across 6 mice, heat map pseudo-color). **c,** Experimental schematic. **d,** SNr section from Npas1-Cre mice after stereotaxic injection of DIO-EYFP (*left*) or DIO-ChR2-EYFP (*middle,right*) into the SNr. Group average is ChR2-EYFP expression across 8 mice. **e-f,** *In vivo* recordings from the SNr showing responses of neurons during an opto-tagging protocol (5 ms width, 20 Hz, 1 mW blue light). The distribution of responses was significantly different in the medial (n = 84 neurons, N = 6 mice) vs. lateral SNr (n = 101 neurons, N = 6 mice); Chi Square Test: χ^2^ (2, 185) = 9.6, p = 0.008, with more neurons excited in the medial vs. lateral SNr (Fisher exact test, p = 0.01). **g-h,** *In vivo* recordings from the SNr showing responses of neurons during a different opto-tagging protocol (1 s width, 0.2 Hz, 1 mW red light). The distribution of responses was similar across medial (n = 52 neurons, N = 6 mice) and lateral territories (n = 87 neurons, N = 6 mice): Fisher Exact test (1, 139), p = 1.0.

To test the hypothesis that excitation of medial SNr neurons promotes persistent behavioral rescue, we sought a strategy to target these neurons directly for optogenetic experiments. Most SNr neurons express PV (Fig. 1b), with expression tending to be higher in lateral SNr neurons compared to medial SNr neurons^4,7^. We asked whether the Npas1-Cre line, which labels non-PV+ cells in the GPe^47^, might also label non/weak-PV+ cells in the SNr, permitting preferential targeting of the medial SNr (Fig. 5c). Injections of DIO-EYFP into the SNr of Npas1-Cre mice did indeed drive preferential expression of EYFP in the medial SNr, with only sparse labeling observed in the rest of the nucleus (Fig. 5d). This pattern was also replicated with injections of DIO-ChR2-EYFP (Fig. 5d).

To confirm that ChR2 expression in Npas1-Cre mice preferentially excites medial SNr neurons, we performed *in vivo* opto-tagging experiments (5 ms pulses delivered at 20 Hz for 5 s, 1 mW) using awake, head-restrained mice. Recordings from medial trajectories within the SNr showed higher proportions of excited neurons than recordings from lateral trajectories (40% medial vs. 23% lateral) (Fig. 5e-f). By comparison, the vast majority of SNr neurons across both medial and lateral territories were inhibited in mice injected with pan-neuronal Jaws (hSyn-Jaws) (1 s pulses delivered every 5 s, 20 trials, 1 mW) (Fig. 1g-h).

### Medial SNr Excitation Promotes Long-Lasting Behavioral Rescue

With the ability to restrict ChR2 expression to the medial SNr, we next sought to assess the effects of stimulating these neurons on movement in dopamine depleted mice (N = 12).

Initially, mice showed no response to medial SNr stimulation, as evidenced by the lack of a light-locked response during the lightON periods (Fig. 6a-b). But over the course of the treatment period, mice gradually became less immobile, and this response persisted for hours after stimulation (Fig. 6a-d). The median therapeutic duration was 2.5 ± 1.4 hrs and half of the mice were still moving at the end of the 3.2 hrs post-stimulation period (Fig. 6d). Immobility was not decreased in control mice (N = 4) that received light stimulation but lacked ChR2 expression (Fig. 6a,c).

**Figure 6:**
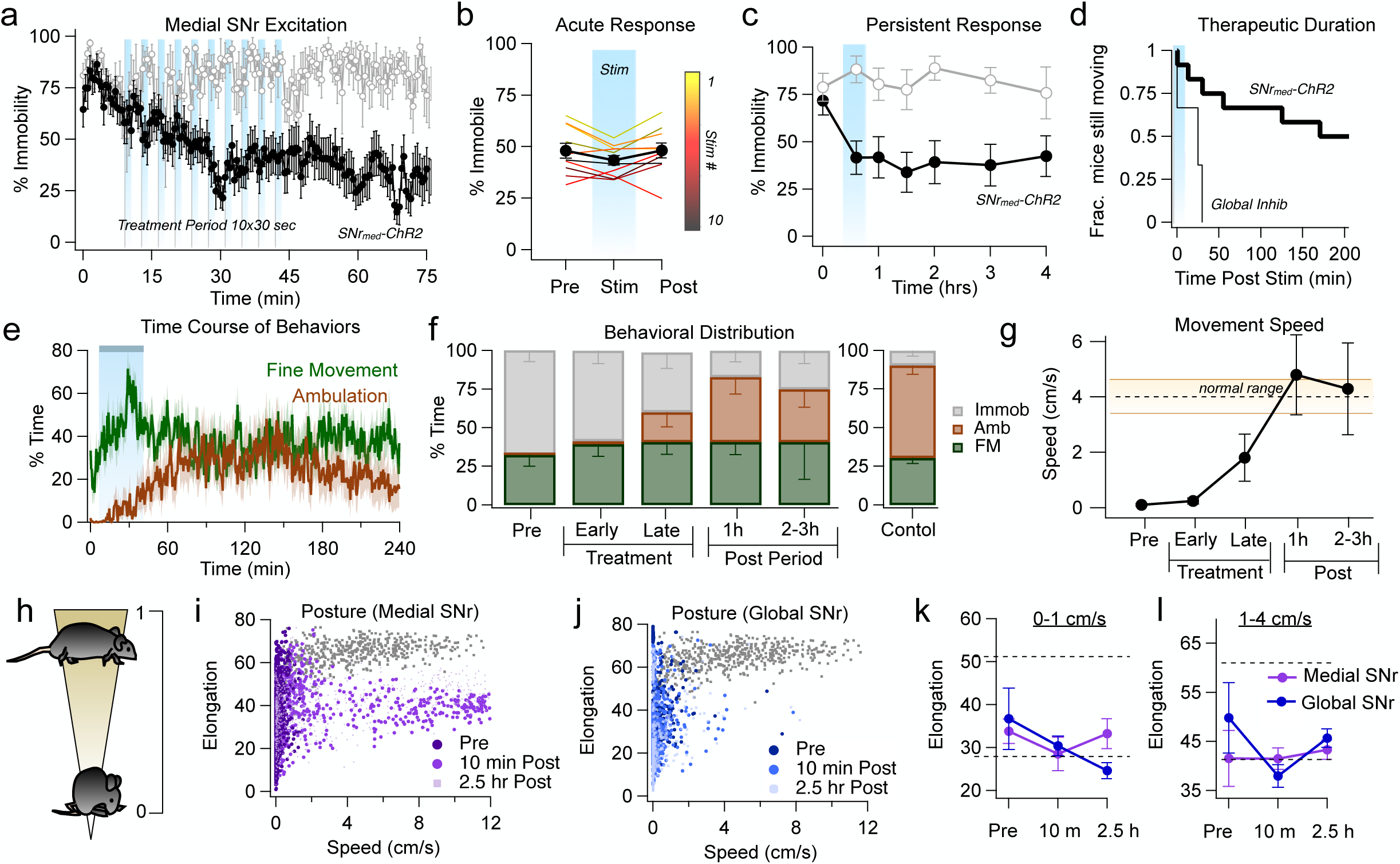
Stimulation of medial SNr promotes movement persistently. **a,** Immobility (percentage of time mice exhibited >0.01% pixel change in 30 s bins) as a function of time during treatment with medial SNr stimulation, (black, N = 12 mice) or in control mice that received light stimulation but lacked viral expression (grey, N = 4 mice). Error bars, sem. Blue lines denote periods of optical stimulation of ChR2: 30 s blue light (1 mW, continuous), spaced 3 min apart. **b,** Acute response to medial SNr stimulation (N = 12 mice). Averages for each trial are shown (1-10, thin lines) as well as the average across all trials (black circles, error bars, sem). Data are from 30 s immediately before stimulation (*pre*), 30 s during stimulation (*stim*), and 30-60 s after each stimulation (*post*). There was no significant change in immobility during the lightON period. Paired t-test: Pre vs. Stim (N = 12 mice) p = 0.34.**c,** Immobility in 10 min bins shows that immobility is persistently reduced by treatment with medial SNr stimulation (N = 12) but not in No virus control mice (N = 4). Repeated measures ANOVA (ChR2 vs no virus): F(1,196) = 14.2, p =0.0002. **d,** Therapeutic duration, calculated from the time of last stimulation until mice returned to within 80% of their initial immobility (*black*). The therapeutic duration from mice treated with global SNr inhibition is re-plotted for reference (*grey*). **e**, Percentage of time mice showing a therapeutic response (N = 8) spent engaged in fine movements (*green*) or ambulation (*brown*) during the experiment. Treatment period is indicated with blue box. **f,** Fraction of time responsive mice spent immobile (*grey*), engaged in fine movements (*green*), or ambulation (*brown*) (N = 8 mice). Behavioral distribution from dopamine intact control mice is plotted for reference (N = 2 mice). Error bars, sem. *Pre*, 10 min. baseline before treatment; *Early*, first 20 min of treatment period; *Late*, Last 20 min of treatment period; *1h*, 10 min period sampled 1 hr after optogenetic treatment; *2-3h*, average of two 10 min bins, sampled 2 hrs and 3 hrs after optogenetic treatment. **g,** Average movement speed measured at the experimental time points described in *f*. Repeated measures ANOVA (N = 8 mice): F(4,28) = 5.09, p 0.003. **h,** Schematic of postural quantification. **i-j,** Body elongation (5 s bins) plotted as a function of movement speed before and after treatment with medial SNr stimulation (N = 8 mice) (*i)* or global SNr inhibition (N = 6 mice) (*j*). Data from dopamine intact controls (grey, Fig. 1) is re-plotted for reference. ***k-l*,** Effect of treatment on body elongation in dopamine depleted mice at low movement speeds (k, 0-1 cm/s) and high movement speeds (l, 1-4 cm/s). Horizontal dashed line denotes average body elongation in control mice over the respective speed ranges, 51.6 ± 4.7 (low speed); 61.0 ± 3.6 (high speed). There was no significant effect of treatment on body posture at either speed and no significant group interaction. 2-Way ANOVA, 0-1 cm/s: SNr_med_, p = 0.70; SNr_global_, p = 0.28; interaction p = 0.30. 2-Way ANOVA, 1-4 cm/s: SNr_med_, p = 0.49; SNr_global_, p = 0.40; interaction p = 0.39.

To examine the nature of behavioral recovery induced by medial SNr stimulation, we analyzed the behavioral patterns of mice in which the therapeutic duration persisted for at least 1 hr after stimulation (8 out of 12 mice). Calculating the percentage of time that mice spent performing “fine movement”, “ambulation”, or “immobile” behaviors revealed that behavioral recovery consisted of a combination of fine motor and ambulatory behaviors (Fig. 6e-f).

Ambulation, a locomotor behavior that was not recovered with global SNr inhibition, became apparent towards the end of the treatment period and persisted throughout the post-treatment period (Fig. 6e-f). After rescue, mice spent ∼80% of their time engaged in movements (a ∼2.4-fold increase over baseline), with nearly equal amounts of time spent performing ambulation or fine movements (Fig. 6e-f). Movement speed in the post-treatment period matched levels seen in dopamine intact mice (Fig. 6g).

Analysis of body elongation revealed that the therapeutic effects of medial SNr stimulation were specific for locomotion. The hunched posture seen in mice after dopamine depletion was not improved by medial SNr stimulation (Fig. 6h-i,k). A graph of body elongation as a function of motor speed shows that mice treated with medial SNr stimulation recover a full range of locomotor speeds, but with markedly less body elongation compared to dopamine intact controls (Fig. 6i-l). Hunched postures were also seen in mice treated with global SNr inhibition, but these mice recovered neither posture nor movement in the post-treatment period (Fig. 6j-l).

### Medial SNr Excitation Persistently Reduces Parkinsonian Pathophysiology

To test whether long-lasting behavioral rescue induced by medial SNr stimulation was accompanied by long-lasting changes in neural activity within the SNr, indicative of network plasticity, we performed recordings from awake head fixed mice (N = 6) before and up to 2 hrs after treatment with medial SNr stimulation (1 mW continuous stim for 30 s, repeated every 3 min). During treatment, 5 out of 10 medial SNr units responded with increases in firing rates that were sustained across the 30 s light pulse and evoked consistently across all 10 stimulations (Fig. 7a-b). The remaining 5 units showed no significant change in firing rate during light the light pulses (Fig. 7a-b).

**Figure 7:**
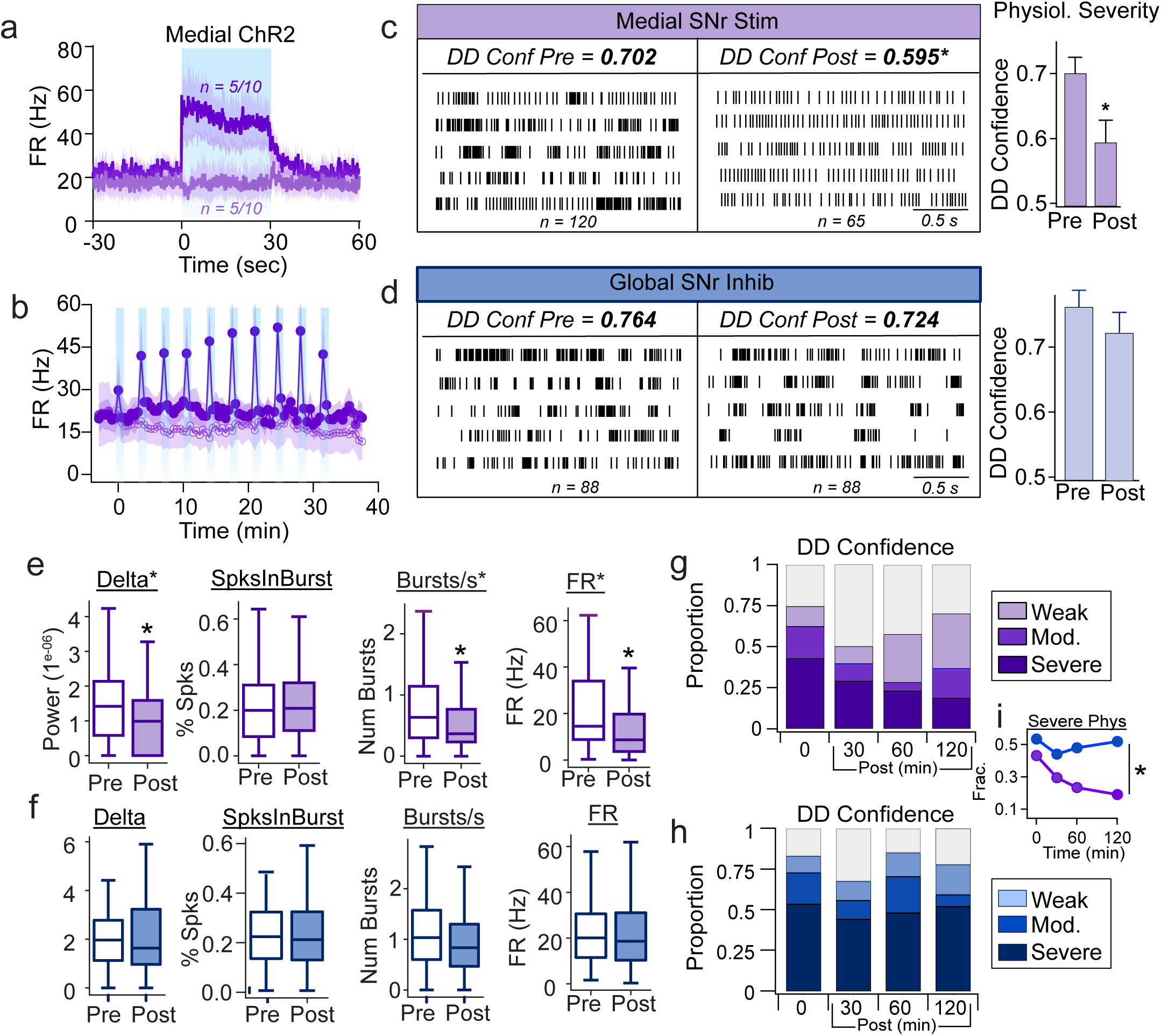
Electrophysiological changes induced by medial SNr excitation are persistent. **a,** Responses of SNr neurons during 30 s light activations of ChR2 (1 mW, continuous blue light, 10 repetitions spaced 3 min apart). Of the 10 neurons where recordings were stable across all 10 stimuli (N = 4 mice), 5 were excited (FR_pre_ 22.8 ± 4.4 Hz vs. FR_stim_ 47.1 ± 11.3 Hz; paired t-test, p = 0.09) and 5 showed no significant change in firing rate (FR_pre_ 17.2 ± 6.5 Hz vs. FR_stim_ 16.6 ± 6.7 Hz; paired t-test, p = 0.42). **b,** Responses were reproducible across all 10 stimuli. **c-d**, Rasters showing spontaneous activity of representative SNr neurons before and after optogenetic treatment with medial SNr stimulation (n_pre_ = 120 neurons, n_post_ = 65 neurons) (*c*) or global SNr inhibition (n_pre_ = 88 neurons, n_post_ = 88 neurons) (*d*). MLP confidence that neurons exhibited parkinsonian pathophysiology was significantly decreased after medial SNr treatment (unpaired t-test, p = 0.01) but not after global inhibition treatment (unpaired t-test, p = 0.37). **e-f,** Persistent effects of optogenetic treatment on electrophysiological parameters. Data are from neurons recorded before treatment (*pre*) and 0-2 hrs after treatment (*post*). Medial SNr stimulation (*e*) drove a significant decrease in delta power (Pre: 1.7e^-06^ ± 1.5e^-06^ vs. Post: 0.99e^-06^ ± 1.1e^-06^; MWU, p= 0.0005), bursts/s (Pre: 0.81 ± 0.66 burst/s vs. Post: 0.52 ± 0.55 bursts/s; MWU, p = 0.0004), and firing rate (Pre: 22.1 ± 18.6 Hz vs. Post: 13.8 ± 13.1 Hz; MWU, p = 0.001); there was no significant change in the fraction of spikes occurring within bursts (Pre: 0.22 ± 0.16 vs. Post: 0.20 ± 0.14; paired t-test, p = 0.41). Global SNr inhibition *(f)* did not drive significant changes in any of these electrophysiological parameters during the 2 hr post-treatment window (Delta Power: p = 0.81; SpikesInBurst: p = 0.88; Bursts/s: p = 0.18; Firing Rate: p = 0.63). **g-h**, The fraction of neurons showing severe parkinsonian pathophysiology (MLP confidence score, >0.83), moderate pathophysiology (MLP confidence score, 0.67-0.83), weak pathophysiology (MLP confidence score, 0.5-0.67) or no pathophysiology (MLP confidence score, < 0.5, grey). A Chi Square Test showed a significant shift in categorical severity after treatment with medial SNr stimulation: χ^2^ (2, 185) = 9.6, p = 0.036; n_pre_ = 120, n_30_ = 27, n_60_ = 17, n_90-120_ = 21, but not after treatment with global SNr inhibition: χ^2^ (2, 185) = 9.6. p = 0.302; n_pre_ = 88, n_30_ = 34, n_60_ = 27, n_90-120_ = 27. **i,** The fraction of neurons classified as having ‘severe’ parkinsonian pathophysiology was significantly decreased across all time points following medial SNr treatment, but not global SNr treatment. 2-Way ANOVA medial vs. global: F(1,7) = 15.9, p = 0.028.

After delivering optogenetic treatment, neural activity in the SNr was monitored for an additional 2 hrs, moving the recording electrode around to different locations to sample activity broadly throughout the nucleus (Fig. 7c). Data collected from these extended recordings sessions were compared against data collected from extended recordings sessions in mice (N = 6) who had received treatment with global SNr inhibition (Fig. 7d).

To assess the effects of treatment on SNr pathophysiology, we entered the electrophysiological features of each neuron into our MLP model and asked the model to report a confidence score that a given unit was from a dopamine depleted animal. Treatment with medial SNr stimulation drove a significant decrease in the MLP network’s confidence scores (Fig. 7c, *p <= 0.05), indicating a decrease in pathophysiological severity. Conversely, global SNr inhibition had no significant effect on confidence scores (Fig. 7d, p = 0.2).

To examine which electrophysiological features were treated by medial SNr stimulation, we first assessed the effects of treatment on delta power and the fraction of spikes in bursts, features that the MLP model relied on heavily to discriminate activity between control and dopamine depleted mice (Fig. 7e-f). In the 2 hrs post period following treatment, there was a significant reduction in the delta power of spike trains seen in mice treated with medial SNr stimulation but not in mice treated with global SNr inhibition (Fig. 7e-f). Neither medial SNr stimulation nor global SNr inhibition reduced the fraction of spikes occurring within bursts (Fig. 7e-f). Medial SNr stimulation also drove persistent decreases in SNr firing rates and number of bursts, whereas global SNr inhibition did not (Fig. 7e-f).

To break these results down further and assess how long physiological changes persisted post-treatment, neurons were classified into four electrophysiological categories: Neurons that received confidence scores of >0.83 for DD pathophysiology were considered to have “severe” pathophysiology; neurons with confidence scores of 0.68 – 0.83 had “moderate” pathophysiology; neurons with confidence scores of 0.5 – 0.67 had “weak” pathophysiology; and neurons with confidence scores <0.5 were considered to be electrophysiologically normal. As shown in Fig. 7g-h, the fraction of SNr units exhibiting “severe” pathophysiology was reduced for the entire 2 hr period following medial SNr stimulation, but not following global SNr inhibition. Although we were not able to collect a robust physiological data set beyond the 2 hr time point due to deteriorating recording quality, neural activity appeared to drift back towards its pre-treatment pathophysiological state, suggesting that suppression of electrophysiological signatures of dopamine depletion persists for ∼ 2 hrs. We saw no significant change in the distribution of units’ pathophysiology levels over time post-stimulation in mice treated with global SNr inhibition (Fig. 7h). Taken together, these results show that medial SNr stimulation reduces pathophysiology in the SNr for ∼ 2 hrs after treatment whereas global inhibition does not.

## Discussion

Our results demonstrate that the spatial pattern of neuromodulation in the SNr determines the time course and duration of locomotor recovery. Global inhibition of the SNr facilitates acute locomotor recovery, enabling movements to occur during periods of active neuromodulation but not beyond. In contrast, focal stimulation restricted to the medial SNr induces a more gradual recovery of movement that is not locked to stimulus periods and persists for hours beyond the treatment period. These results suggest the presence of parallel, independent pathways that mediate acute vs. persistent locomotor responses. Persistent effects at the behavioral level were mirrored by persistent effects at the cellular level, with medial SNr stimulation attenuating parkinsonian pathophysiology for hours beyond treatment. These results prescribe a spatial pattern of neuromodulation in the SNr to achieve sustained behavioral recovery and promote network plasticity in a mouse model of advanced dopamine depletion.

### Biologically-inspired treatments for improved therapeutic outcomes

Stimulation of the basal ganglia has been used for decades to treat the motor symptoms of Parkinson’s disease. Conventional DBS, which consists of continuous, high frequency stimulation, is thought to mediate its effects by disrupting aberrant network patterning that develops in the basal ganglia under conditions of low dopamine^9,10,13,14,17,18^. But when stimulation is turned off, motor symptoms and network pathophysiology return within minutes, suggesting that conventional stimulation masks symptoms but does not restore circuit function. The delivery of high frequency electrical stimulation to neural tissue seems to have the capacity to drive plasticity that could alter circuit dynamics long-term. Indeed, evidence suggests that DBS does alter neuronal excitability and synaptic dynamics of circuits both inside and outside the basal ganglia^48–52^. But conventional DBS appears not to access this potential for long-term changes.

Alternative biologically-inspired therapies seek to leverage the plastic properties of the nervous system to train damaged circuits out of their pathological state. An example of a biologically-inspired treatment for Parkinson’s disease is coordinated reset DBS (CR-DBS), a stimulation pattern designed to use spike-timing dependent plasticity to counteract the hyper-synchronized network state that develops in the basal ganglia under conditions of low dopamine^14,53–55^. Early results suggested that CR-DBS provides long-lasting therapeutic benefits that extend for hours, and even days after treatment, but widespread implementation of the technique has been hampered by variability in its efficacy and complex stimulation requirements that cannot be readily implemented with standard medical devices^56–58^.

Adaptive DBS (aDBS) uses knowledge about electrophysiological biomarkers of aberrant basal ganglia activity to control the timing or amplitude of neurostimulation in a closed-loop manner^9,14,15,59,60^. Adaptive DBS to counteract aberrant oscillatory activity in beta or gamma frequency bands has proven effective at delivering the same levels of therapeutic benefits comparable to conventional DBS, but with significantly less overall stimulation time, reducing treatment-related side effects^61–63^. Although aDBS can transiently interrupt aberrant activity patterns, it remains to be seen whether it has long-lasting effects on underlying network pathophysiology.

CR-DBS and aDBS are examples of how biologically-inspired innovations in the timing and pattern of stimulation have led to improved outcomes for patients. Identifying and targeting relevant brain circuits is also critical to maximize therapeutic benefits while minimizing side effects. Computational reconstruction of the neural pathways activated during DBS has shed light on relevant therapeutic networks in a number of diseases such as Parkinson’s disease, OCD, Tourette’s and dystonia^64–67^.

The use of optogenetics can identify cellular targets with which to access these therapeutic networks. In the GPe, for example, cell-type specific manipulation is more therapeutically effective than global stimulation and engages persistent therapeutic effects^19^. The direct use of optogenetics in humans is still not widely available, but insights about the basic biology of the relevant cell populations in the GPe led to the development of a human-translatable electrical stimulation paradigm^20^ that has proven to be safe and effective in patients and requires 80% less stimulation time than conventional DBS^68^. Important questions still remain about the identity of the neural circuits required for long-lasting effects and how to direct and tailor stimulation to ensure therapeutic success.

### Biomarkers of long-lasting therapeutic effects

DBS is typically programmed through a trial-and-error process that relies on immediate behavioral feedback to tune stimulus parameters. Because stimulation that induces long-lasting therapeutic effects can sometimes require 10-20 minutes to take effect^19,20^ (Fig. 6), current practices in DBS programming might actually bias clinicians away from parameters that could provide long-lasting benefits.

Our results identify both short-term and long-term electrophysiological biomarkers that predict whether a given intervention will engage long-lasting therapeutic effects. Acutely, stimulation should drive asymmetrical responses in basal ganglia output nuclei, such that medial territories are preferentially excited with respect to lateral territories. Because all basal ganglia nuclei project to the output nuclei, we anticipate that the desired pattern of activity will be achievable from multiple different stimulation loci throughout the circuit.

Long-term, we identified several biomarkers of network plasticity associated with long-lasting behavioral changes. The most predictive single parameter was a decrease in delta power (0.5-4 Hz). Elevated delta power is commonly detected in the local field potentials (LFP) of Parkinson’s patients^69–71^, but has been understudied because it was believed to be a side effect of anesthesia^72–74^. But increased delta power is present even in awake mice, for which it is one of the best predictors of movement severity and dopamine depletion^26^.

The mechanisms underlying increased delta power are not known, but computational modeling suggests they might be an emergent property of synchronized GABAergic networks oscillating out of phase with one another^75,76^. Decreases in local GABA concentrations within the SNr are predicted to exacerbate delta oscillations through effects on the local inhibitory collateral network^76^. Recently, GABAergic tone in the SNr was found to depend on GABA co-release from dopaminergic neurons in the SNc, and the loss of these inputs is tightly correlated with the onset of motor deficits in the MCI-Park mouse model of Parkinson’s disease^27,77^. The dopamine neurons responsible for GABA co-release in the SNr, ALDH1A1-type dopamine neurons, send the majority of their projections to the medial SNr^78^, similar to the region critical for long-lasting therapeutic effects in our study. A transient intervention applied locally to the medial SNr might be more effective at decoupling activity between medial and lateral territories of the nucleus, enabling the network to transition out of its pathologically synchronized state.

An argument for local plasticity mechanisms within the SNr is further supported by the finding that therapeutic stimulation had only mixed effects on burst firing in the SNr, activity that is likely driven by excitatory inputs from the STN^4^. Medial SNr stimulation reduced the number of bursts occurring in the SNr (consistent with results seen after GPe interventions^19^), but did not reduce the percentage of spikes occurring within bursts, which our MLP classifier suggests is a better predictor of network pathophysiology.

The behavioral effects of SNr interventions in our study were specific for bradykinesia, but not postural symptoms. Both features are cardinal symptoms of Parkinson’s disease, but bradykinesia is hypothesized to result from basal ganglia dysfunction whereas postural symptoms are thought to be driven by circuits outside of the basal ganglia that are less responsive to dopamine replacement therapy. The specificity of our interventions for bradykinesia but not posture suggest that the basal ganglia are the locus of plasticity in our study.

### Neural Pathways for Behavioral Persistence

The finding that stimulation of the medial SNr recovers movement is surprising and counterintuitive to predictive models of basal ganglia function. Typically, DBS is targeted to lateral (sensorimotor) areas of basal ganglia nuclei because these sites provide immediate behavioral effects and are tightly coupled to thalamo-cortical motor circuits^79^. Disinhibition of these canonical motor circuits is the most likely explanation for the acute behavioral recovery induced by optogenetic inhibition of the SNr. But behavioral recovery induced by medial SNr stimulation follows a different time course than the acute effects induced by global SNr inhibition, requiring tens of minutes to take full effect and persisting for hours after stimulation. Dopamine disorders can affect all functional loops of the basal ganglia, not just motor circuits^80^.

Correspondingly, our results suggest that persistent vs. acute therapeutic effects might be driven through dissociable pathways.

Based on topographical organization, the medial SNr is considered to be part of the limbic basal ganglia circuit and its neurons have been shown to play a role in regulating an animal’s arousal state^7,8,81^. The medial SNr also projects directly to the pontine reticular nucleus, oral part (PnO), a pathway important for limb control and turning^82^. Unilateral stimulation of this pathway was sufficient to counteract the asymmetrical rotation phenotype observed in unilaterally dopamine depleted mice^82^.

While there are still limitations to the duration of therapeutic effects induced by medial SNr stimulation, these effects persist for hours longer than conventional intervention strategies. The persistent effects of medial SNr stimulation might be induced through different circuits and different alterations in pathophysiology than are currently impacted by existing DBS therapies and hence suggest alternative targets and strategies to restore behavior. These findings represent important steps towards developing strategies to repair circuit function despite the absence of dopamine and to thereby promote sustained recovery of movement.

## Materials and Methods

### Animals

Experiments were conducted in accordance with the guidelines from the National Institutes of Health and with approval from Carnegie Mellon University Institutional Animal Care and Use Committee. Adult male and female mice (>8 weeks old) on a C57BL/6J background were used for experiments. All mice were maintained on a 12-h light/12-h dark cycle with food and water provided ad libitum. Behavioral, in vivo electrophysiological, and histological experiments involving optogenetic excitation of medial SNr neurons were performed in Npas1-cre transgenic mice. Behavioral experiments involving SNr neuron ablation and SNr cell counting experiments were performed in Pvalb-2A-cre (PV-cre) transgenic mice. Histological experiments involving Lhx6_only_-GPe projections to the SNr were performed in Lhx6-iCre;PV-Flp mice.

### Surgery (virus injections and dopamine depletions)

All surgical procedures were performed using aseptic techniques and were conducted under general anesthesia. AAV vectors were injected into the substantia nigra pars reticulata (SNr) or the external segment of the globus pallidus (GPe) following methods previously described^19^.

Briefly, anesthesia was induced using ketamine (100 mg/kg) and xylazine (30 mg/kg) and maintained throughout surgery using 1.5% isoflurane. Mice were placed in a stereotaxic frame (Kopf Instruments), where the scalp was opened and bilateral holes were drilled in the skull (SNr: -3.2 mm AP, +/-1.45 mm ML from Bregma; GPe: -0.3 mm AP, +/- 2.0 mm ML from Bregma). Viral constructs were injected with a Nanoject (Drummond Scientific) through a pulled glass pipette (tip diameter ∼30 mm) whose tip was positioned below the top of the skull (SNr: -4.75 mm DV, 150 nL; GPe: -3.65 mm DV, 150 nL). To prevent backflow of virus, the pipette was left in the brain for 5 min after completion of the injection. All experiments were performed at least 3-4 weeks after injection to allow time for full viral expression. At that point, some mice underwent a second surgery to make craniotomies and implantation of head-bars for in vivo extracellular recordings under anesthesia as described above. Head-bars were implanted to secure mice for in vivo extracellular recordings. Bilateral craniotomies were created over the SNr: (-3.2 mm AP, +/- 1.5 mm ML) and a stainless steel head-bar was fixed to the skull with Metabond dental cement. Dental cement was extended from the head-bar to surround the extent of both craniotomies to form a well. This well was then filled with silicone elastomer (Kwik-sil, WPI) to prevent infection and damage to the exposed brain tissue. For behavioral experiments, optic fiber cannulae were implanted in the SNr (coordinates described above) using Metabond dental cement. In addition, in the same surgery, mice were dopamine depleted with 6-hydroxydopamine (6-OHDA). Bilateral holes were drilled over the medial forebrain bundle (MFB; -1.3 mm AP, pm1.15 mm lateral from bregma) for 6-OHDA injections. A pulled glass pipette (tip diameter ∼30 µm) attached to a Nanoject was slowly lowered into the MFB (4.95 mm DV) and allowed to settle for 5 min. At this point, 1 µL of 6-OHDA (5 µg/µL) in 0.9% NaCl was slowly injected into the MFB at a rate of 0.1 µL/min. The injection cannula was left in place for 7-10 more min before retracting. Post-surgery, animals were injected with 3mg/kg Ketofen s.c. and allowed to recover on a heating pad before being returned to their standard home cage. After dopamine depletions, mice were individually housed and placed in a recovery station. The station consisted of a new cage, soft food, trail mix, a shallow water dish, and half of the cage was placed on a heating pad. In addition, a daily injection of saline (0.9% NaCl; intraperitoneally) was used to curb dehydration, and weight was closely observed to monitor the health of each animal.

### AAV vectors

Injections of purified double-floxed AAV2-DIO-ChR2-eYFP or AAV2-DIO-ChR2-mCherry (cell-specific medial SNr activation), AAV2-DIO-eYFP (cell-specific medial SNr labeling), AAV5-hSyn-Jaws-KGC-GFP-ER2 (global SNr inhibition), AAV2-Flex-DTR-GFP (SNr cell necrosis), AAV5-EF1a-Nuc-Flox(mCherry)-EGFP (SNr cell counting) or rAAVDJ-pAAV-Ef1a-FRT-DIO-eGFP (Lhx6_only_-GPe projections to the SNr) (B. Lim) were made in the SNr or GPe of Npas1-cre, PV-cre and Lhx6-iCre;PV-Flp transgenic mice (8-14 weeks old).

### Optogenetic stimulation

For all implants, optic fibers (Thorlabs), cut to length and glued inside 1.25mm ceramic ferrules, were matched for light transmission and calibrated to achieve the desired output measured at the tip of optic fiber cannula (PM100D, Thorlabs). For behavioral experiments, excitation of medial SNr (Npas1+) neurons was performed with constant 1mW blue light (465 nm) from high powered LEDs (Optogenetics-Dual-LED, Prizmatix) (N = 12 mice). Global inhibition of the SNr was performed with a constant 1mW red light (630 nm) from high powered LEDs (Ultrahigh-Power LED Controller, Prizmatix) (N = 6 mice). For *in vivo* electrophysiological experiments, excitation of medial SNr (Npas1+) neurons was performed with constant 1mW blue light (465 nm) from high powered LEDs (PlexBright LED 465 nm, Plexon) (N = 6 mice). Global inhibition of the SNr was performed with a constant 1mW red light (630 nm) from high powered LEDs (PlexBright LED 630 nm, Plexon) (N = 6 mice). For behavioral and in vivo electrophysiological experiments, a 30 s long light pulse was delivered in the SNr. This light pulse was repeated ten times, separated by 3 min.

### Behavioral experiments

For opto-manipulation experiments, four to five days after 6-OHDA lesion mice were bilaterally connected to an optogenetic dual fiber (Optogenetics-Dual-LED or Ultrahigh-Power LED Controller, Prizmatix) and placed in the center of a square open field arena (50x50 cm). The optogenetic dual fiber was adjusted to ensure 1 mW of power at the tip of the previously implanted ferrules. Mice activity was monitored from overhead and the side of the arena. The body center point, base of the tail, and nose were tracked using Noldus EthoVision software. After collecting 10 min of baseline spontaneous activity, a 30 s long pulse of light was delivered with the corresponding LED and was repeated ten times, separated by 3 min intervals. Following the final stimulation bout mice were observed for another 3 hrs to examine the persistence of behavioral intervention^19^. For experiments involving SNr neuron ablation, mice injected with DTR in the SNr were placed in the center of a square open field arena (50x50 cm) for three consecutive days to record their baseline spontaneous locomotion. Mice were then injected with diphtheria toxin (DTX) (1µg) (N = 5) or saline (N = 3) intraperitoneally. Two weeks after DTX or saline injection mice were placed in the open field arena again to record a pre 6-OHDA or sham lesion measurement. Finally, four to five days after 6-OHDA or sham lesion mice were placed in the open field arena for the last testing session. Their activity was monitored for 20 min as described above.

### Acclimatation to head fixation and *in vivo* extracellular recordings of SNr neurons

Mice were placed and secured on top a running wheel and allowed to run freely for 45-120 min four days before the recording session. On the day of the recording, mice were fixed and secured on top of the wheel and allowed 15 min to acclimate to the head-fixed position. Then, the silicone elastomer was removed and the craniotomy was cleaned, the well was filled with 0.9% NaCl and used as a ground reference. A linear 16-channel silicon opto-probe with sites spaced 25 µm apart (A1x16-10mm-25-177-OA16LP, fiber diameter: 105 core and 0.66 numerical aperture, Neuronexus) was attached to the micromanipulator and centered on Bregma. The opto-probe was slowly advanced (5–7µm/s) until the top of the recording target. Once a population of responsive units was identified, the spontaneous firing activity of these putative SNr units was recorded. During the pre and post opto-stimulation periods, once the firing activity of a set of SNr neurons was recorded the opto-probe was advanced to record a new set of neurons. Due to the scale of our craniotomy, when the opto-probe was advanced to the bottom of the SNr (∼5.00 mm), the opto-probe was taken out of the brain and reinserted in a different coordinate. This allowed sampling across the SNr. During the full optogenetic stimulation protocol (see: Optogenetic stimulation), the opto-probe was left at the same recording site for the full duration, including an additional 5-30 min after stimulation.

### Optical tagging

In medial SNr activation electrophysiological experiments, an opto-tagging strategy was used to classify SNr neurons as positive or negative for channelrhodopsin (ChR2)^19^. Brief pulses of blue light (5 ms width, 20Hz) were applied at the end of each recording session before advancing the opto-probe to a new location. Peristimulus histograms (PSTH) were created (bin size = 1 ms) to analyze the responsiveness of the recorded neurons to the opto-tagging protocol. Baseline activity (10 ms pre stimulation) for each light pulse was compared to the activity observed in the post pulse period (40 ms post stimulation). Neurons were considered putative ChR2+ when a significant increase in their activity was detected within 0-10 ms of light onset. Neurons inhibited or not responding to the opto-tagging protocol were considered putative ChR2-. Similarly, in global SNr inhibition (hSyn-Jaws) electrophysiological experiments, longer (1 s width, 0.2Hz) red light pulses were applied to determine the proportion of SNr neurons directly inhibited by the light. Peristimulus histograms (PSTH) were created (bin size = 50 ms) to analyze the responsiveness of the recorded neurons to the opto-tagging protocol. Baseline activity (1 s pre stimulation) for each light pulse was compared to the activity observed during the light pulse (1 s light ON). Neurons were considered inhibited when a significant decrease in their activity was detected within the light pulse.

### Spike sorting

Data was sampled and stored at 32 kHz and filtered at 150–8000 Hz for spiking activity using an OmniPlex amplifier (Plexon, Inc.). Spike detection, clustering and sorting were performed using Offline Sorter (Plexon). For classification as a single unit, the following criteria were set: the principal component analysis of waveforms generated a cluster of spikes significantly distinct from other unit or noise clusters (P < 0.05), the J3-statistic was > 1, and the Davies–Bouldin statistic was < 0.5. Finally, the sorted spike trains were manually inspected on the basis of spike waveform, stability across time and inter-spike interval refractory period violations (< 0.35% of interspike intervals (ISIs) were < 2 ms). In the case where a unit was lost during recording, it was only used in analysis for the time period when its spike cluster satisfied these criteria, and only if its cluster was present for at least 3 min^26^.

### Histology

Mice were anesthetized with a lethal dose of ketamine (100 mg/kg) and xylazine (30 mg/kg) and perfused transcardially with phosphate buffered saline (PBS), followed by 4% paraformaldehyde (PFA) in PBS. Brains were retrieved and post-fixed in 4% PFA for 24-hrs before being placed in a 30% sucrose solution. Brains were sliced in 30 µm thick coronal sections using a freezing microtome (Microm HM 430; Thermo Scientific). Brain sections were stored in a cryoprotectant (30% glycerol, 40% PBS, 30% ethylene glycol) solution until processed for histological validation. For SNr ablation experiments, SNr sections of mice injected with AAV2-Flex-DTR-GFP were incubated with a rabbit anti-GFP (1:1000, Millipore Sigma, AB3080) primary antibody to enhance the GFP signal at 4 °C overnight. The primary antibody was then detected with an anti-rabbit Alexa-Fluor488 (1:500, Fischer Scientific, A21206) secondary antibody, incubated at room temperature for 3 hrs. Then, tissue was mounted and cover-slipped with Vectashield antifade mounting medium (Vector). For SNr cell counting experiments, SNr sections of mice injected with AAV5-EF1a-Nuc-Flox(mCherry)-EGFP were mounted and cover-slipped with Vectashield antifade mounting medium (Vector). For histological experiments characterizing the projection pattern of Lhx6_only_-GPe neurons in the SNr, GPe and SNr sections of mice injected with rAAVDJ-pAAV-Ef1a-FRT-DIO-eGFP were incubated with a rabbit anti-GFP (1:1000, Millipore Sigma, AB3080) primary antibody to enhance the GFP signal at 4 °C overnight. The primary antibody was then detected with an anti-rabbit Alexa-Fluor488 (1:500, Fischer Scientific, A21206) secondary antibody, incubated at room temperature for 3 hrs. Then, tissue was mounted and cover-slipped with Vectashield antifade mounting medium (Vector). All mounted tissue was imaged and scanned at 10x resolution using a fluorescence microscope (BZ-X series, Keyence). For behavioral and *in vivo* electrophysiological experiments, virus expression and placement of optic fibers and recording probe were also histologically validated. To confirm virus expression and the position of optic fibers and recording probes used for behavioral and in vivo electrophysiological experiments, brain sections containing the SNr were mounted and cover-slipped with Vectashield antifade mounting medium (Vector). Mounted tissue was then imaged and scanned at 10x resolution using a fluorescence microscope (BZ-X series, Keyence). Virus expression in the SNr was visually confirmed in each mouse in the rostro-caudal and the medio-lateral axis. The location of optic fibers implanted in the SNr was determined based on tissue damage around the tip of the optic fiber. Electrode tracks created by the recording probe were used to reconstruct the proper location of the probe within the SNr. Animals were excluded from the study when optic fiber or recording probe locations were not found or found outside of the target area or virus expression was inadequate.

### Tyrosine hydroxylase quantification

The degree of dopamine denervation was assessed in all mice based on the immunofluorescence against tyrosine hydroxylase (TH) as described previously^83^. Slices containing the striatum were incubated in a rabbit anti-TH (1:500, Pel-Freeze, P40101-150) primary antibody at 4°C for 24 hrs. The primary antibody was then detected with an Alexa-Fluor 647-conjugated donkey anti-rabbit (1:500, Fisher Scientific, A31573) secondary antibody, incubated at room temperature for 90 min. Then, tissue was mounted and cover-slipped with Vectashield antifade mounting medium (Vector). Sections containing the dorsal striatum were bilaterally imaged using an epifluorescence microscope at 10× magnification (BZ-X series, Keyence). To analyze the fluorescence intensity, we used the pixel intensity-measuring tool in ImageJ. A 100 × 100 µm square from each hemisphere was measured (background fluorescence was subtracted by measuring fluorescence intensity in cortex) and normalized to the pixel intensities measured in littermate control mice tissue that was processed and imaged in parallel. All mice included in this study had <20% TH remaining on both brain hemispheres.

### Data analysis

*Behavioral experiments.* Mouse behavior was analyzed by top-down camera (29.97 FPS). Positions of nose, base of the tail, and body center of each mouse were tracked using EthoVision XT software (Noldus). In optogenetic experiments, ambulation bouts were scored post hoc as periods of 2.40 cm/s movement (lasting for >0.5 s and >0.5 s apart). Immobility bouts were scored by using a criterion of >0.01% pixel change in 0.5 s, lasting for >0.5 s and >0.5 s apart. Fine movement bouts were scored as periods where the mouse was not ambulating and not immobile lasting for >0.5 s and >0.5 s apart. Spontaneous locomotion was measured via distance traveled (cm/bin) measurements. In SNr ablation experiments, velocity (cm/s), distance traveled (cm/bin) and elongation (Noldus elongation metric; 30 sample moving average; average/bin) were measured.

Elongation (E) was computed as follows:

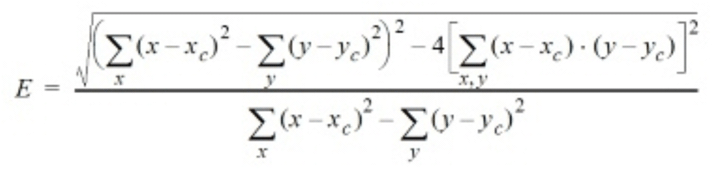

where x and y are pixel coordinates of the animal’s outline and x_c_ and y_c_ mark the animal’s center point. The resulting value is a percentage value where 0% corresponds to a circular body shape and 100% corresponds to a linear shape.

#### In vivo electrophysiological experiments

Spike trains of sorted individual SNr units were analyzed through Neuroexplorer in which firing rate and coefficient of variation were calculated in 30 s bins. To analyze the effects of the opto-stimulation protocols on SNr neurons’ firing rate, PSTHs (bin size: 100 ms) were generated using Spike2 CED. The PSTHs to analyze the effects of opto-tagging protocols on SNr neurons’ firing activity were generated using Spike2 CED (see: Optical tagging). Further data analysis was performed using a Multi-Layer Perceptron classifier constructed in Python (see Computational neuron classification section).

#### Histological experiments

##### SNr ablations in PV-cre mice injected with DTR-GFP

SNr ablations were validated by comparing the expression of enhanced GFP fluorescence signal in the SNr in mice injected with DTX to the expression in mice injected with saline. SNr cell counting. PV-cre mice (N = 2) bilaterally injected with color changing reporter virus Nuc-Flox(mCherry)-eGFP in the SNr were used to measure the proportion of SNr neurons expressing PV. This viral construct expresses the fluorescent reporter eGFP in the presence of cre while it expresses mCherry in the absence of cre. SNr neurons expressing eGFP or mCherry were counted using the FIJI Cell counter tool (ImageJ, NIH) in 12 SNr sections.

##### Lhx6_only_-GPe projections to the SNr

Lhx6-iCre;PV-Flp mice (N = 6) injected with the intersectional virus rAAVDJ-pAAV-Ef1a-FRT-DIO-eGFP in the GPe were used to determine Lhx6_only_-GPe neuron axonal projection patterns to the SNr. SNr sections (approximate Bregma - 3.52) across mice were aligned using the BigWarp plugin in ImageJ and an average image was generated using the concatenate and z-stack ImageJ built-in functions. The average image showing cell-specific targeting of medial SNr neurons in Npas1-cre mice (N = 8) was generated using the same procedure.

### Computational neuron classification

#### Classifier structure and training

We developed a Multi-Layer Perceptron (MLP) neural network to classify recordings as coming from SNr neurons in dopamine intact or dopamine depleted animals. The network architecture, built using Scikit-Learn in Python, consisted of four layers: an input layer, two hidden layers with 200 and 100 units respectively, and an output layer. All layers used the ReLU activation function and were connected in a feed-forward manner. The input layer received a normalized feature vector, which was passed to the first hidden layer, where each node computed a weighted sum of inputs to which it applied the ReLU activation function. This process was repeated in the second hidden layer. The output layer aggregated the activations from the second hidden layer using a weighted sum and applied a sigmoid function to produce a probability score, or confidence, for classification of the spike train as either control or depleted.

The training data comprised spike train features computed from 30 s recordings of 512 units from control animals (N = 14 mice) and 623 units from depleted animals (N = 26 mice). The features used to train the network included: firing rate, coefficient of variation (CV), delta power, beta power, bursts per second, average burst firing rate, percent time spent bursting, percent of spikes within bursts, average burst duration, average interburst interval, non-bursting firing rate, and burst-related firing rate increase. Detailed definitions of these features are provided in Table 1 and Table 2.

**Table 1:** Table of notation for spike train and burst properties.

| Property | Symbol | Property | Symbol |
| --- | --- | --- | --- |
| number of bursts | $N_b$ | $j$th spike time in $i$th burst | $T_i^j$ |
| $i$th burst duration | $D_i$ | final spike time in $i$th burst | $T_i^{-1}$ |
| $i$th burst firing rate | $f_i$ | number of spikes in $i$th burst | $N_i$ |
| spike train recording length | $L$ | overall spike train firing rate | $f$ |
| number of spikes in spike train | $N$ | mean of feature $f$ | $\langle f \rangle$ |

**Table 2:**
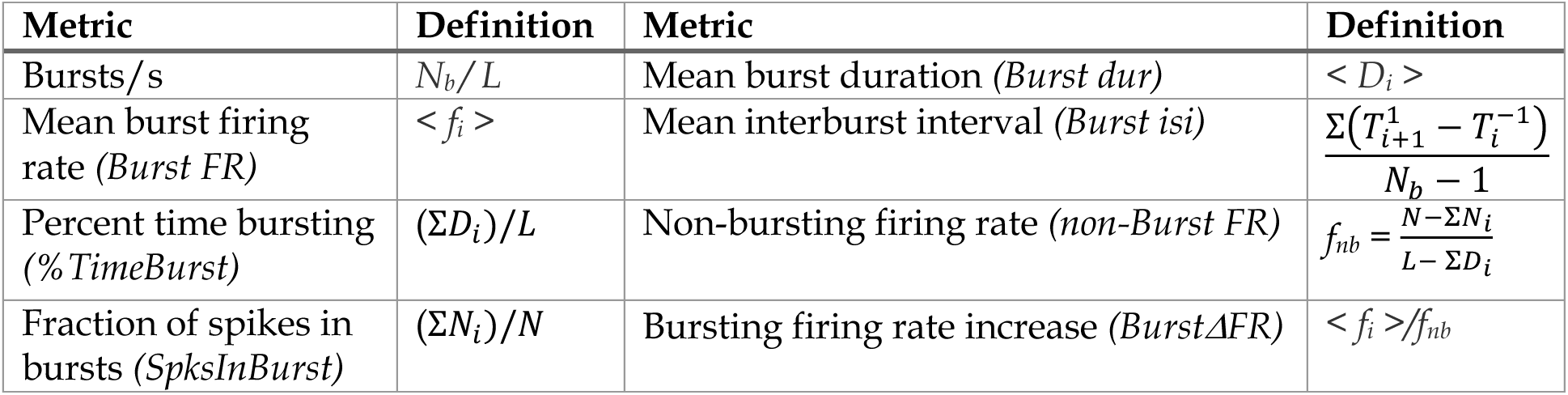
Definitions of bursting features using the notation from Table 1. All sums are taken from *i=1* to *i=N_b_* except the upper limit of summation for mean interburst interval is *N_b_-1*.

Before presentation to the classifier, data was preprocessed to remove outliers. For this step, features within control and DD data sets were z-scored separately. Outliers were identified as any spike train that contained at least one feature with a z-score magnitude greater than 3, leaving n = 462 control units and n = 551 DD units after outlier removal. The data set was split into training (80%) and testing (20%) subsets, and this splitting was repeated 15 times with different random seeds to train 15 distinct MLP models using back-propagation. Each training subset was normalized, and the corresponding testing subset was normalized to the training subset. Training accuracy averaged 79%, with precision scores of 75% for control units and 84% for DD units, and recall scores of 81% for control units and 78% for DD units. Testing accuracy averaged 71%, with precision scores of 67% for control units and 75% for DD units, and recall scores of 71% for both control and DD units. The confidence score was defined as the probability that a unit was obtained from depleted conditions, as provided by the MLP model.

#### Burst metrics

Detection of bursts was performed using the Poisson surprise method^84^. The reference period for burst likelihood was twice the spike train maximum inter-spike interval preceding the burst candidate, rather than being based on the overall spike train as originally defined. The surprise threshold was S = 3, the minimum number of spikes for the burst was set at 3, and the maximum number of spikes for a burst was set at 10.

#### Code

The code repository for spike train analysis and further implementation details is hosted on GitHub (https://github.com/jparker25/Aristieta_Parker_Rubin_Gittis_2024_motor_rescue) and will be made publicly available upon acceptance.

## References

1. Nelson, A. B. & Kreitzer, A. C. Reassessing models of basal ganglia function and dysfunction. Annu Rev Neurosci 37, 117–135 (2014).

2. McGregor, M. M. & Nelson, A. B. Circuit Mechanisms of Parkinson’s Disease. Neuron 101, 1042–1056 (2019).

3. Galvan, A. & Wichmann, T. Pathophysiology of Parkinsonism. Clinical Neurophysiology vol. 119 1459–1474 (2008).

4. Delgado-Zabalza, L. et al. Targeting parvalbumin-expressing neurons in the substantia nigra pars reticulata restores motor function in parkinsonian mice. Cell Rep. 42, 113287 (2023).

5. Willard, A. M. et al. State transitions in the substantia nigra reticulata predict the onset of motor deficits in models of progressive dopamine depletion in mice. Elife 8, (2019).

6. Wichmann, T. et al. Comparison of MPTP-induced changes in spontaneous neuronal discharge in the internal pallidal segment and in the substantia nigra pars reticulata in primates. Exp Brain Res 125, 397–409 (1999).

7. McElvain, L. E. et al. Specific populations of basal ganglia output neurons target distinct brain stem areas while collateralizing throughout the diencephalon. Neuron 109, 1721– 1738 (2021).

8. Liu, D. et al. A common hub for sleep and motor control in the substantia nigra. Science (80-.). 367, 440–445 (2020).

9. Gittis, A. H. & Sillitoe, R. V. Circuit-Specific Deep Brain Stimulation Provides Insights into Movement Control. Annu. Rev. Neurosci. 47, (2024).

10. Perlmutter, J. S. & Mink, J. W. Deep brain stimulation. Annu. Rev. Neurosci. 29, 229–257 (2006).

11. Schor, J. S. et al. Therapeutic deep brain stimulation disrupts movement-related subthalamic nucleus activity in parkinsonian mice. Elife 11, e75253 (2022).

12. Zhuang, Q.-X. et al. Regularizing firing patterns of rat subthalamic neurons ameliorates parkinsonian motor deficits. J. Clin. Invest. (2018) doi:10.1172/JCI99986.

13. Jakobs, M., Fomenko, A., Lozano, A. M. & Kiening, K. L. Cellular, molecular, and clinical mechanisms of action of deep brain stimulation—a systematic review on established indications and outlook on future developments. EMBO Mol. Med. 11, e9575 (2019).

14. Krauss, J. K. et al. Technology of deep brain stimulation: current status and future directions. Nat. Rev. Neurol. 17, 75–87 (2021).

15. Neumann, W.-J., Horn, A. & Kühn, A. A. Insights and opportunities for deep brain stimulation as a brain circuit intervention. Trends Neurosci. 46, 472–487 (2023).

16. Temperli, P. et al. How do parkinsonian signs return after discontinuation of subthalamic DBS? Neurology 60, 78–81 (2003).

17. Rubin, J. E. & Terman, D. High frequency stimulation of the subthalamic nucleus eliminates pathological thalamic rhythmicity in a computational model. J. Comput. Neurosci. 16, 211–235 (2004).

18. Rosenbaum, R. et al. Axonal and synaptic failure suppress the transfer of firing rate oscillations, synchrony and information during high frequency deep brain stimulation. Neurobiol. Dis. 62, 86–99 (2014).

19. Mastro, K. J. et al. Cell-specific pallidal intervention induces long-lasting motor recovery in dopamine-depleted mice. Nat. Neurosci. 20, 815–823 (2017).

20. Spix, T. A. et al. Population-specific neuromodulation prolongs therapeutic benefits of deep brain stimulation. Science 374, 201–206 (2021).

21. Mastro, K. J., Bouchard, R. S., Holt, H. A. K. & Gittis, A. H. Transgenic mouse lines subdivide external segment of the globus pallidus (GPe) neurons and reveal distinct GPe output pathways. J. Neurosci. 34, 2087–2099 (2014).

22. Albin, R. L., Young, A. B. & Penney, J. B. The functional anatomy of basal ganglia disorders. Trends Neurosci 12, 366–375 (1989).

23. DeLong, M. R. Primate models of movement disorders of basal ganglia origin. Trends in Neurosciences vol. 13 281–285 (1990).

24. Okun, M. S. & Vitek, J. L. Lesion therapy for Parkinson’s disease and other movement disorders: update and controversies. Mov. Disord. 19, 375–389 (2004).

25. Puelles, L., Stühmer, T., Rubenstein, J. L. R. & Diaz, C. Critical test of the assumption that the hypothalamic entopeduncular nucleus of rodents is homologous with the primate internal pallidum. J. Comp. Neurol. 531, 1715–1750 (2023).

26. Whalen, T. C., Willard, A. M., Rubin, J. E. & Gittis, A. H. Delta oscillations are a robust biomarker of dopamine depletion severity and motor dysfunction in awake mice. J. Neurophysiol. 124, 312–329 (2020).

27. González-Rodríguez, P. et al. Disruption of mitochondrial complex I induces progressive parkinsonism. Nature 599, 650–656 (2021).

28. Deumens, R., Blokland, A. & Prickaerts, J. Modeling Parkinson’s disease in rats: an evaluation of 6-OHDA lesions of the nigrostriatal pathway. Exp Neurol 175, 303–317 (2002).

29. Azim, E., Jiang, J., Alstermark, B. & Jessell, T. M. Skilled reaching relies on a V2a propriospinal internal copy circuit. Nature 508, 357–363 (2014).

30. Madisen, L. et al. A toolbox of Cre-dependent optogenetic transgenic mice for light-induced activation and silencing. Nat. Neurosci. 15, 793–802 (2012).

31. Sethi, K. Levodopa unresponsive symptoms in Parkinson disease. Mov. Disord. 23 Suppl 3, S521–33 (2008).

32. Takakusaki, K., Saitoh, K., Harada, H. & Kashiwayanagi, M. Role of basal ganglia-brainstem pathways in the control of motor behaviors. Neurosci. Res. 50, 137–151 (2004).

33. Takakusaki, K., Habaguchi, T., Ohtinata-Sugimoto, J., Saitoh, K. & Sakamoto, T. Basal ganglia efferents to the brainstem centers controlling postural muscle tone and locomotion: a new concept for understanding motor disorders in basal ganglia dysfunction. Neuroscience 119, 293–308 (2003).

34. Aristieta, A., Ruiz-Ortega, J. A., Miguelez, C., Morera-Herreras, T. & Ugedo, L. Chronic L-DOPA administration increases the firing rate but does not reverse enhanced slow frequency oscillatory activity and synchronization in substantia nigra pars reticulata neurons from 6-hydroxydopamine-lesioned rats. Neurobiol. Dis. 89, 88–100 (2016).

35. Wichmann, T. et al. Comparison of MPTP-induced changes in spontaneous neuronal discharge in the internal pallidal segment and in the substantia nigra pars reticulata in primates. Exp. Brain Res. 125, 397–409 (1999).

36. Wichmann, T. & Soares, J. Neuronal firing before and after burst discharges in the monkey basal ganglia is predictably patterned in the normal state and altered in parkinsonism. J Neurophysiol 95, 2120–2133 (2006).

37. Wang, Y. et al. Changes in firing rate and pattern of GABAergic neurons in subregions of the substantia nigra pars reticulata in rat models of Parkinson’s disease. Brain Res. 1324, 54–63 (2010).

38. Seeger-Armbruster, S. & von Ameln-Mayerhofer, A. Short- and long-term unilateral 6-hydroxydopamine lesions in rats show different changes in characteristics of spontaneous firing of substantia nigra pars reticulata neurons. Exp Brain Res 224, 15–24 (2013).

39. Lobb, C. J., Zaheer, A. K., Smith, Y. & Jaeger, D. In vivo electrophysiology of nigral and thalamic neurons in alpha-synuclein-overexpressing mice highlights differences from toxin-based models of parkinsonism. J Neurophysiol 110, 2792–2805 (2013).

40. Kravitz, A. V et al. Regulation of parkinsonian motor behaviours by optogenetic control of basal ganglia circuitry. Nature 466, 622–626 (2010).

41. Gerfen, C. R. et al. D1 and D2 dopamine receptor-regulated gene expression of striatonigral and striatopallidal neurons. Science (80-.). 250, 1429–1432 (1990).

42. Assaf, F. & Schiller, Y. A chemogenetic approach for treating experimental Parkinson’s disease. Mov. Disord. 34, 469–479 (2019).

43. Lozano, A. M. & Lang, A. E. Pallidotomy for Parkinson’s Disease. Neurosurg. Clin. N. Am. 9, 325–336 (1998).

44. Cif, L. & Hariz, M. Seventy Years with the Globus Pallidus: Pallidal Surgery for Movement Disorders Between 1947 and 2017. Mov. Disord. 32, 972–982 (2017).

45. Aristieta, A., Parker, J. E., Gao, Y. E., Rubin, J. E. & Gittis, A. H. Dopamine depletion weakens direct pathway modulation of SNr neurons. Neurobiol. Dis. 196, 106512 (2024).

46. Dodson, P. D. et al. Distinct developmental origins manifest in the specialized encoding of movement by adult neurons of the external globus pallidus. Neuron 86, 501–513 (2015).

47. Hernández, V. M. et al. Parvalbumin+ Neurons and Npas1+ Neurons Are Distinct Neuron Classes in the Mouse External Globus Pallidus. J. Neurosci. Off. J. Soc. Neurosci. 35, 11830–11847 (2015).

48. Ni, Z. et al. Pallidal deep brain stimulation modulates cortical excitability and plasticity. Ann. Neurol. 83, 352–362 (2018).

49. Milosevic, L. et al. Modulation of inhibitory plasticity in basal ganglia output nuclei of patients with Parkinson’s disease. Neurobiol. Dis. 124, 46–56 (2019).

50. Milosevic, L. et al. Neuronal inhibition and synaptic plasticity of basal ganglia neurons in Parkinson’s disease. Brain 141, 177–190 (2018).

51. Steiner, L. A. et al. Persistent synaptic inhibition of the subthalamic nucleus by high frequency stimulation. Brain Stimul. 15, 1223–1232 (2022).

52. Neumann, W.-J., Steiner, L. A. & Milosevic, L. Neurophysiological mechanisms of deep brain stimulation across spatiotemporal resolutions. Brain 146, 4456–4468 (2023).

53. Tass, P. A. et al. Coordinated reset has sustained aftereffects in Parkinsonian monkeys. Ann Neurol 72, 816–820 (2012).

54. Ebert, M., Hauptmann, C. & Tass, P. A. Coordinated reset stimulation in a large-scale model of the STN-GPe circuit. Front. Comput. Neurosci. 8, 154 (2014).

55. Adamchic, I. et al. Coordinated reset neuromodulation for Parkinson’s disease: proof-of-concept study. Mov Disord 29, 1679–1684 (2014).

56. Wang, J. et al. Coordinated Reset Deep Brain Stimulation of Subthalamic Nucleus Produces Long-Lasting, Dose-Dependent Motor Improvements in the 1-Methyl-4-phenyl-1,2,3,6-tetrahydropyridine Non-Human Primate Model of Parkinsonism. Brain Stimul. 9, 609–617 (2016).

57. Kromer, J. A., Khaledi-Nasab, A. & Tass, P. A. Impact of number of stimulation sites on long-lasting desynchronization effects of coordinated reset stimulation. Chaos An Interdiscip. J. Nonlinear Sci. 30, 83134 (2020).

58. Wang, J. et al. Shuffling Improves the Acute and Carryover Effect of Subthalamic Coordinated Reset Deep Brain Stimulation. Front. Neurol. 13, 716046 (2022).

59. Meidahl, A. C. et al. Adaptive Deep Brain Stimulation for Movement Disorders: The Long Road to Clinical Therapy. Mov. Disord. 32, 810–819 (2017).

60. Gilron, R. et al. Long-term wireless streaming of neural recordings for circuit discovery and adaptive stimulation in individuals with Parkinson’s disease. Nat. Biotechnol. 39, 1078–1085 (2021).

61. Oehrn, C. R. et al. Chronic adaptive deep brain stimulation versus conventional stimulation in Parkinson’s disease: a blinded randomized feasibility trial. Nat. Med. (2024) doi:10.1038/s41591-024-03196-z.

62. Little, S. et al. Adaptive deep brain stimulation in advanced Parkinson disease. Ann. Neurol. 74, 449–457 (2013).

63. Tinkhauser, G. et al. The modulatory effect of adaptive deep brain stimulation on beta bursts in Parkinson’s disease. Brain 140, 1053–1067 (2017).

64. Hollunder, B. et al. Mapping dysfunctional circuits in the frontal cortex using deep brain stimulation. Nat. Neurosci. 27, 573–586 (2024).

65. Rajamani, N. et al. Deep brain stimulation of symptom-specific networks in Parkinson’s disease. Nat. Commun. 15, 4662 (2024).

66. Hollunder, B. et al. Toward personalized medicine in connectomic deep brain stimulation. Prog. Neurobiol. 210, 102211 (2022).

67. Horn, A. et al. Deep brain stimulation induced normalization of the human functional connectome in Parkinson’s disease. Brain 142, 3129–3143 (2019).

68. Kariv, S. et al. Pilot Study of Acute Behavioral Effects of Pallidal Burst Stimulation in Parkinson’s Disease. Mov. Disord. 39, 1873–1877 (2024).

69. Du, G. et al. Properties of oscillatory neuronal activity in the basal ganglia and thalamus in patients with Parkinson’s disease. Transl. Neurodegener. 7, 17 (2018).

70. Levy, R. et al. Dependence of subthalamic nucleus oscillations on movement and dopamine in Parkinson’s disease. Brain A J. Neurol. 125, 1196–1209 (2002).

71. Zhuang, P., Hallett, M., Meng, D., Zhang, Y. & Li, Y. Characteristics of oscillatory activity in the globus pallidus internus in patients with Parkinson’s disease (P1. 8–028). Neurology 92, P1-8 (2019).

72. Ruskin, D. N., Bergstrom, D. A. & Walters, J. R. Nigrostriatal lesion and dopamine agonists affect firing patterns of rodent entopeduncular nucleus neurons. J. Neurophysiol. 88, 487–496 (2002).

73. Belluscio, M. A., Kasanetz, F., Riquelme, L. A. & Murer, M. G. Spreading of slow cortical rhythms to the basal ganglia output nuclei in rats with nigrostriatal lesions. Eur. J. Neurosci. 17, 1046–1052 (2003).

74. Walters, J. R., Hu, D., Itoga, C. A., Parr-Brownlie, L. C. & Bergstrom, D. A. Phase relationships support a role for coordinated activity in the indirect pathway in organizing slow oscillations in basal ganglia output after loss of dopamine. Neuroscience 144, 762–776 (2007).

75. Whalen, T. C., Parker, J. E., Gittis, A. H. & Rubin, J. E. Transmission of delta band (0.5-4 Hz) oscillations from the globus pallidus to the substantia nigra pars reticulata in dopamine depletion. J. Comput. Neurosci. (2023) doi:10.1007/s10827-023-00853-z.

76. Phillips, R. S., Rosner, I., Gittis, A. H. & Rubin, J. E. The effects of chloride dynamics on substantia nigra pars reticulata responses to pallidal and striatal inputs. Elife 9, e55592 (2020).

77. Simmons, D., Ding, J., Khakh, B., Awatramani, R. & Surmeier, D. J. Tonic dendritic GABA release by substantia nigra dopaminergic neurons. bioRxiv (2024) doi:10.1101/2024.03.27.586699.

78. Simmons, D., Ding, J., Khakh, B., Awatramani, R. & Surmeier, D. J. Tonic dendritic GABA release by substantia nigra dopaminergic neurons. bioRxiv (2024) doi:10.1101/2024.03.27.586699.

79. Wichmann, T. & DeLong, M. R. Deep-brain stimulation for basal ganglia disorders. Basal Ganglia vol. 1 65–77 (2011).

80. Cavallo, A. & Neumann, W.-J. Dopaminergic reinforcement in the motor system: Implications for Parkinson’s disease and deep brain stimulation. Eur. J. Neurosci. 59, 457– 472 (2024).

81. Foster, N. N. et al. The mouse cortico–basal ganglia–thalamic network. Nature 598, 188– 194 (2021).

82. Cregg, J. M., Sidhu, S. K., Leiras, R. & Kiehn, O. Basal ganglia-spinal cord pathway that commands locomotor gait asymmetries in mice. Nat. Neurosci. 27, 716–727 (2024).

83. Willard, A. M. M., Bouchard, R. S. S. & Gittis, A. H. H. Differential degradation of motor deficits during gradual dopamine depletion with 6-hydroxydopamine in mice. Neuroscience 301, 254–267 (2015).

84. Legendy, C. R. & Salcman, M. Bursts and recurrences of bursts in the spike trains of spontaneously active striate cortex neurons. J. Neurophysiol. 53, 926–939 (1985).

